# Stage-dependent tau post-translational modifications map the spatiotemporal progression of Alzheimer’s disease

**DOI:** 10.64898/2026.04.10.717615

**Authors:** Axelle A.T. Vanparys, Clémence Balty, Manuel Johanns, Nathalie Kyalu Ngoie Zola, Gaëtan Herinckx, Marine Van Calsteren, Nuria Suelves, John L. Woodard, Didier Vertommen, Pascal Kienlen-Campard, Bernard J. Hanseeuw

## Abstract

Alzheimer’s disease (AD) is defined by progressive tau aggregation, yet the molecular events driving this process remain poorly understood. Post-translational modifications (PTMs) are key regulators of tau biology and potential biomarkers of disease progression. Using immunoprecipitation-mass spectrometry and absolute quantification of tau isoforms, we profiled tau PTMs in soluble and insoluble brain fractions. We studied multiple brain regions (hippocampus, inferior temporal and frontal gyri), representing regions affected at different stages of pathology, from human donors spanning the AD spectrum and staged by ABC neuropathological scoring. We uncovered a stage-dependent PTM landscape across AD progression: early phosphorylation changes, including pT217 and pS262, precede later ubiquitination events, such as uK311, associated with tau aggregation. We also identified PTMs negatively correlated with aggregation, including mK258, suggesting potential protective roles. These findings refine our understanding of the spatiotemporal evolution of tau biochemistry and offer mechanistic and translational insights into AD tauopathy.

## INTRODUCTION

Aging is known as the primary driver of Alzheimer’s disease (AD)^1^. As global life expectancy increases, the prevalence of age-related diseases like AD continues to rise, representing a major societal and economic burden worldwide. AD is the leading cause of dementia, accounting for 60 to 70% of cases ^2^. The number of people living with AD was around 57 million in 2019 and is expected to increase to 153 million by 2050 ^3,4^. This dramatic increase highlights the urgent need to unravel the molecular mechanisms underlying AD pathogenesis and to develop effective disease-modifying therapies. Although new biomarkers are emerging for AD ^5,6^, early diagnosis remains challenging in clinical practice ^7,8^. This limitation is critical as immunotherapies targeting amyloid-β (Aβ) pathology are now becoming available, offering new therapeutic perspectives for AD patients ^9,10^.

AD is defined as a secondary tauopathy ^11,12^, characterized by the extracellular deposition of Aβ peptides and intracellular aggregation of hyperphosphorylated tau protein, resulting in the formation of neurofibrillary tangles (NFTs) ^2,13–15^. Aβ plaque formation is widely considered to be the starting point of AD ^14,16^, followed by tau hyperphosphorylation and aggregation, which impair tau’s physiological functions ^2,12,13,17^. To date, neuropathological examination of post-mortem brain tissue is still the gold standard to stage AD progression or to distinguish between tauopathies. Since 2012, AD-related neuropathological changes are classified using the three following parameters: (i) Aβ plaque score (derived from the Thal phase) ^17–19^, (ii) NFTs stage (according to Braak stages) ^13,18^ and (iii) neuritic plaque score (derived from the CERAD score, reflecting the concentration of plaques containing both Aβ and tau aggregates) ^17,18,20^. These three criteria are at the base of the “ABC” scoring system, which categorizes the level of AD neuropathological change as: (i) not, (ii) low, (iii) intermediate, or (iv) high, reflecting the likelihood that dementia is attributable to AD neuropathological hallmarks. The ABC score, therefore, reflects the regional distribution of the previously described neuropathological hallmarks of AD ^17^.

The distribution of tau aggregates is closely correlated with the clinical manifestations of AD and the extent of neurodegeneration ^13,21^, whereas the stages of Aβ deposition are more loosely associated with cognitive impairment ^19^. Moreover, promising biomarker research is currently focused on tau ^5–7^, positioning this protein as target for AD research. Being a “natively unfolded” protein, tau is greatly influenced by posttranslational modifications (PTMs) ^22^. Approximately one-third of the tau sequence can be modified. The clustering of PTMs within distinct motifs and functional domains strongly suggests that tau function is regulated by a complex interplay among these modifications ^22^. Phosphorylation has long been associated with tau aggregates found in tauopathies, and PTMs-driven models of tau aggregation have been proposed ^12,22–26^. What remains elusive is which PTMs drive tau self-association in specific contexts, and which directly assist its physiological function and are trapped as bystanders in tau aggregates.

Tau pathology likely begins with alterations in tau phosphorylation on serine(S)/threonine(T) residues, induced by external factors, triggering or facilitating the appearance of PTMs on other residues ^26^. Lysine residues (K) are of particular interest, as they can undergo multiple competing PTMs, including methylation, acetylation and ubiquitination, all of which have been found in tau ^25,26^. Among these, ubiquitination deserves particular attention, as it appears to underline distinct molecular NFT conformations and modifications specific to each tauopathy ^25,27,28^. However, the spatiotemporal sequence of phosphorylation, ubiquitination and other modifications has not been thoroughly studied during AD progression.

In this study, we used liquid chromatography–coupled tandem mass spectrometry (LC-MS/MS) to decipher how the tau PTM profile changes during AD progression. A human cohort (N = 16) spanning different AD stages (i.e., different ABC scores) was studied. We analyzed Sarkosyl-soluble and -insoluble fractions from three different brain regions hippocampus (HIPP), inferior temporal gyrus (ITG), and inferior frontal gyrus (IFG), which are differentially affected by tau pathology during AD progression. We identified PTMs that positively or negatively correlate with tau aggregation, highlighting their potential mechanistic role in tau deposition. Furthermore, as soluble tau species are more likely to be released into and detected in body fluids ^29^, our findings in the soluble fraction may facilitate the identification of novel biomarkers to stage AD progression in the brain.

## RESULTS

### Subjects and brain samples characteristics

This study used human post-mortem brain tissue obtained from the Netherlands Brain Bank (NBB) and the UCLouvain brain donation program. In total, 16 subjects spanning different stages of AD progression were included. Subjects were classified according to their ABC score (defined by Thal phase, Braak stage and CERAD score, as described in Hyman et al. ^17^), grouping them according to their level of AD neuropathological change (low, intermediate and high). These groups, hereafter referred to as “AD score”, reflect the extent of neuropathological alterations independently of clinical dementia status.

The main characteristics of the cohort are provided in Table1 and Supp Table1. Clinically, subjects were either non-demented controls or diagnosed with Alzheimer’s disease dementia. Neuropathologically, the cohort spanned the full AD spectrum, ranging from Braak stage I to VI and Thal phases 0-5. No significant differences were found between groups for sex, brain weight, or post-mortem delay. Subject groups differed significantly in age at death, with individuals in the “high AD neuropathological score” group, being younger (p < 0.01). As expected, AD neuropathological staging parameters differed significantly across groups, with severe pathology in the high-score group (A3, B3, C3), moderate pathology in the intermediate group (mainly A2–A3, B2, C1–C2), and minimal changes in the low-score group (A0–A1, B1–B2, C0). Consistently, the frequency of clinical AD dementia increased across groups (0/5, 3/7, and 4/4).

**Table 1.**
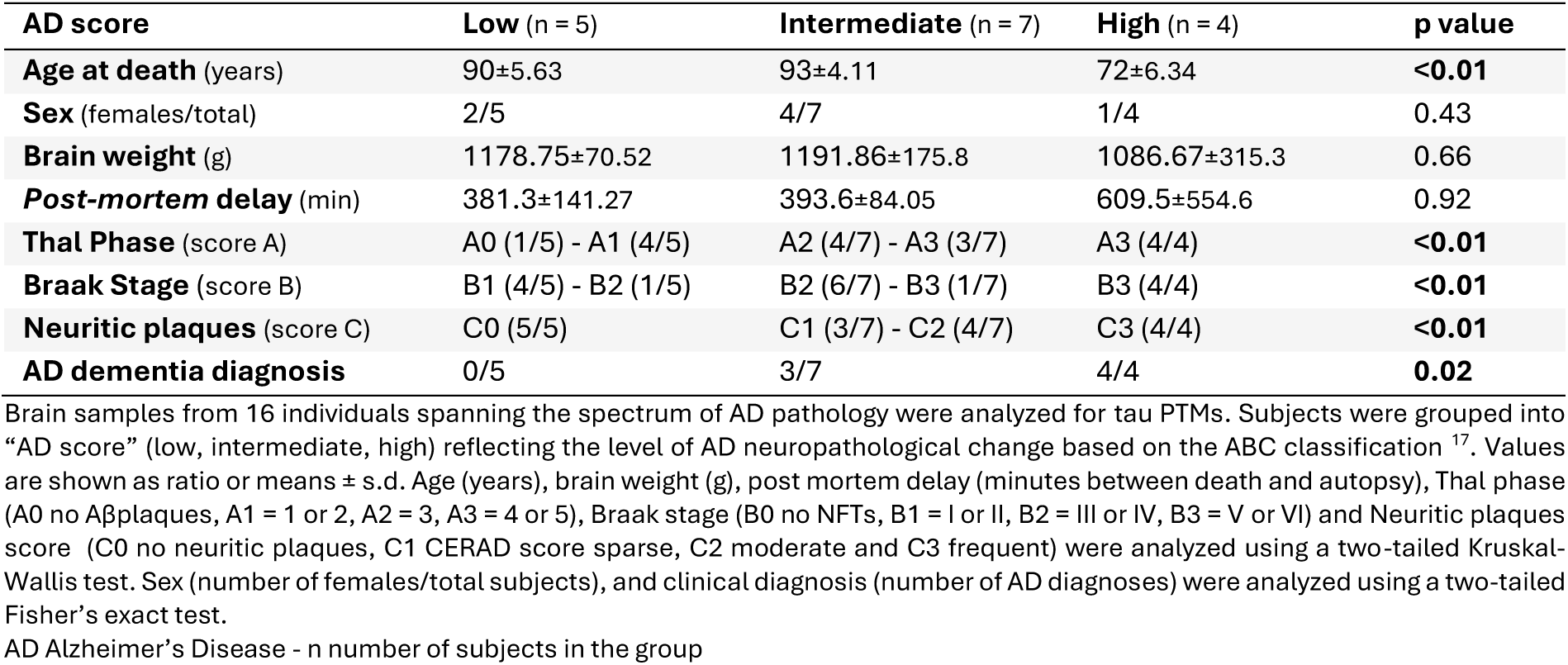
Characteristics of the cohort.

| **AD score** | **Low (n = 5)** | **Intermediate (n = 7)** | **High (n = 4)** | **p value** |
| --- | --- | --- | --- | --- |
| **Age at death** (years) | 90±5.63 | 93±4.11 | 72±6.34 | **<0.01** |
| **Sex** (females/total) | 2/5 | 4/7 | 1/4 | 0.43 |
| **Brain weight** (g) | 1178.75±70.52 | 1191.86±175.8 | 1086.67±315.3 | 0.66 |
| **Post-mortem delay** (min) | 381.3±141.27 | 393.6±84.05 | 609.5±554.6 | 0.92 |
| **Thal Phase** (score A) | A0 (1/5) - A1 (4/5) | A2 (4/7) - A3 (3/7) | A3 (4/4) | **<0.01** |
| **Braak Stage** (score B) | B1 (4/5) - B2 (1/5) | B2 (6/7) - B3 (1/7) | B3 (4/4) | **<0.01** |
| **Neuritic plaques** (score C) | C0 (5/5) | C1 (3/7) - C2 (4/7) | C3 (4/4) | **<0.01** |
| **AD dementia diagnosis** | 0/5 | 3/7 | 4/4 | **0.02** |

Brain samples from 16 individuals spanning the spectrum of AD pathology were analyzed for tau PTMs. Subjects were grouped into “AD score” (low, intermediate, high) reflecting the level of AD neuropathological change based on the ABC classification<sup>17</sup>. Values are shown as ratio or means ± s.d. Age (years), brain weight (g), post mortem delay (minutes between death and autopsy), Thal phase (A0 no Aβ plaques, A1 = 1 or 2, A2 = 3, A3 = 4 or 5), Braak stage (B0 no NFTs, B1 = I or II, B2 = III or IV, B3 = V or VI) and Neuritic plaques score (C0 no neuritic plaques, C1 CERAD score sparse, C2 moderate and C3 frequent) were analyzed using a two-tailed Kruskal-Wallis test. Sex (number of females/total subjects), and clinical diagnosis (number of AD diagnoses) were analyzed using a two-tailed Fisher’s exact test.
AD Alzheimer’s Disease - n number of subjects in the group

For all subsequent analyses, the previously described “AD score” will be used to define sample groups. To model the stereotypical spatiotemporal progression of tau pathology, three brain regions were examined: the hippocampus (HIPP), affected at early stages (Braak stages I-II); the Inferior Temporal Gyrus (ITG), affected at “intermediate” stages (Braak stages III-IV); and the Inferior Frontal Gyrus (IFG), affected at late stages (Braak stages V-VI).

### Absolute quantification of tau aggregates by LC-MS recapitulates neuropathological AD staging

To validate the reported unequal representation of tau protein domains within pathological aggregates (i.e. the enrichment of peptides containing the microtubule-binding region in the insoluble fraction of AD brains ^25,30^) and to assess its evolution across AD progression in different brain regions, we performed absolute quantification of tau domains and isoforms using SureQuant MS analysis. Sarkosyl-insoluble tau was analyzed as a proxy for aggregated tau, as this fraction is widely used to isolate and study aggregated tau species from brain tissue, including paired helical filaments and other fibrillar structures ^31^.Quantification relied on heavy-labeled peptides specific to either N-terminal inserts (N isoforms), the mid domain, and the repeat (R) domain (3R and 4R isoforms). Total microtubule binding region (MTBR) abundance was calculated by summing the 3R and 4R isoforms.

The R isoforms were strongly overrepresented in aggregates compared to the other tau domains across AD scores (Supp Fig.1a). A bigger proportion of the 4R domain was observed compared to the 3R domain in all our conditions (Supp Fig.1a). General insoluble tau accumulation was subsequently assessed based on the MTBR domain (3R+4R) as a proxy. An accumulation of MTBR in aggregates was observed with AD progression, starting in the HIPP of participants classified as “low” stage (Fig.1). This accumulation was also detected in the ITG and HIPP of individuals at the “intermediate” stage. Finally, the increase in MTBR was observed across all the studied regions in the individuals at the “high” stage. In this group, no significant differences were observed between regions, suggesting a plateau in tau accumulation in the more advanced AD stages. We observed a similar increase in the other tau domains (Supp Fig.1b). Together, these findings indicate that the transition from low to intermediate AD stages is characterized by the emergence of insoluble tau deposition in the ITG, consistent with the stereotypical regional progression of tau pathology.

**Fig. 1.**
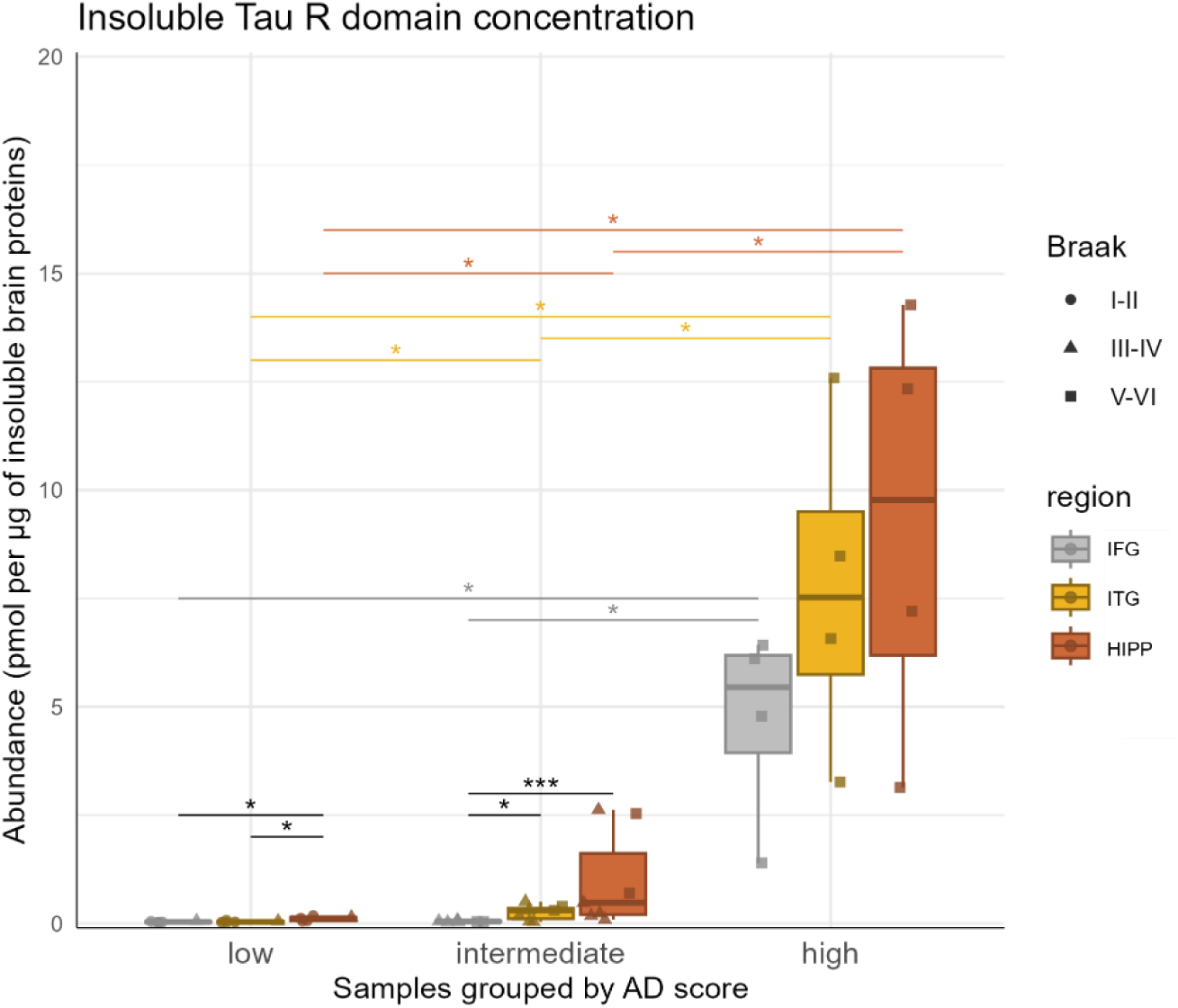
Insoluble tau MTBR concentration during AD progression across several brain regions. Tau MTBR domain (3R+4R) concentration from insoluble protein fraction was measured using mass spectrometry (SureQuant method) in three brain regions (IFG Inferior Frontal Gyrus – ITG Inferior Temporal Gyrus – HIPP Hippocampus) from 16 patients grouped by AD score (low: Low AD score, int: Intermediate AD score, high: High AD score, reflecting the level of AD neuropathological change based on the ABC classification^17^). Statistical comparisons were made between AD scores for each brain region using the Wilcoxon rank-sum test after a Kruskal-Wallis test and between brain regions from patients with a given AD score using a Conover test after a Friedman test. Significant BH-adjusted p-values are indicated in black for the comparison between regions or in the color of the region for comparisons between AD scores: * > 0.05; ** > 0.01; *** > 0.001 MTBR Microtubule binding region – AD Alzheimer’s Disease

### Insoluble tau PTMs act as biochemical markers of AD stages

Sarkosyl-insoluble proteins were analyzed by LC-MS/MS to identify tau PTMs, specifically S and T phosphorylation (p) and K methylation (m), acetylation (a) and ubiquitination (u). A coverage of 98% of the sequence of the full-length tau protein (2N4R) was obtained in the insoluble fraction, ensuring comprehensive representation of all major functional domains in subsequent analyses. We identified 48 individual PTMs in insoluble brain fractions across the entire cohort, including canonical AD-associated phosphorylation sites such as pS202, pT205 and pT404 ^21,32^. The normalized abundance of each modification was compared across the three AD scores, within specific brain regions, to identify PTMs associated with AD progression. We also carried out a regional analysis, comparing the abundance of each

PTM between brain regions at a given AD score (Table2 & Supp Table2), to outline PTMs that increase or decrease during AD progression across brain regions. The abundance of 41 tau PTMs was significantly altered during AD progression and brain regions (Table 2 & Supp Table 2).

**Table 2.**
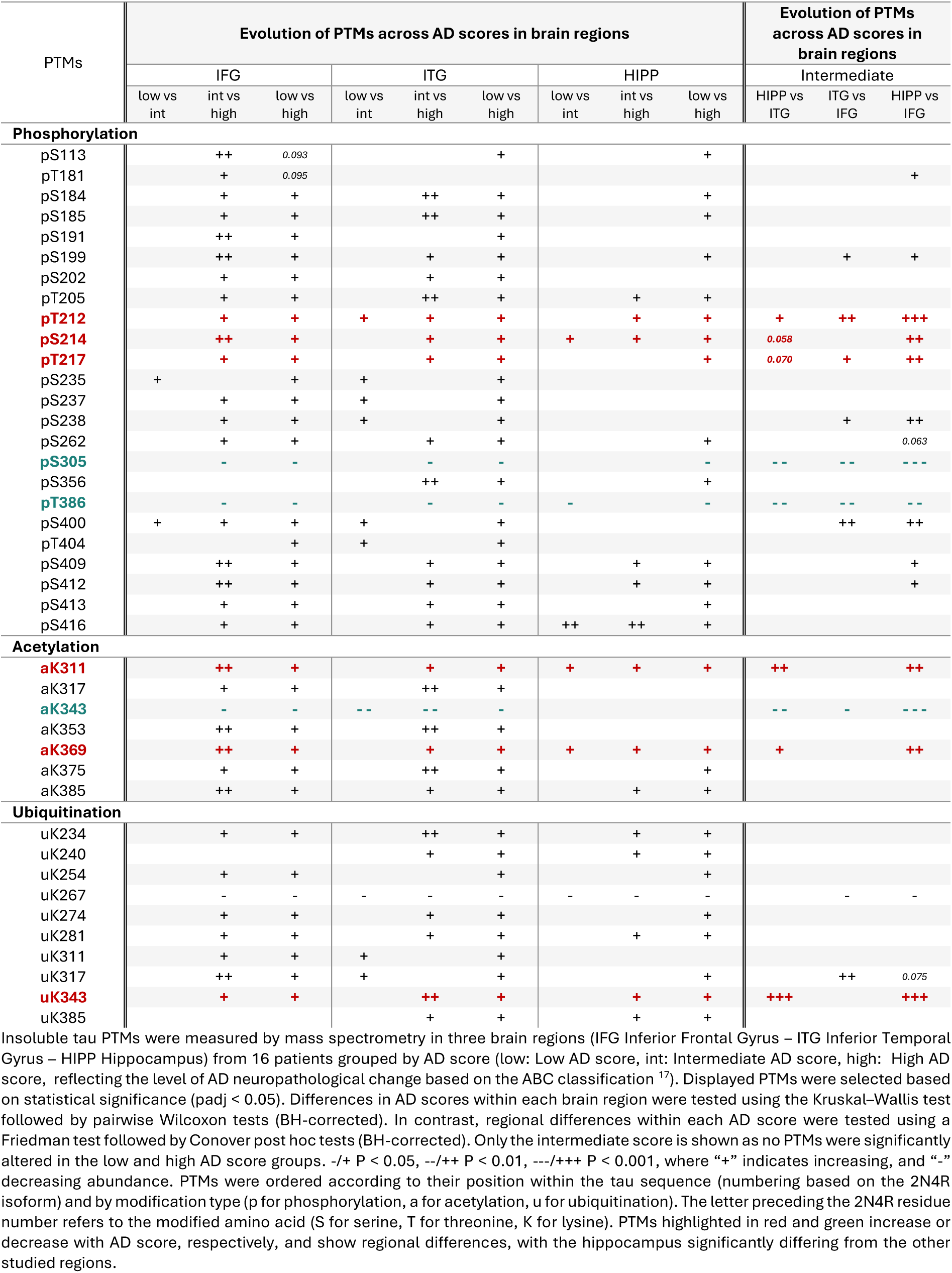
Insoluble tau PTMs altered during AD progression across brain regions and between AD score.

Numerous tau phosphorylation (24), acetylation (7), and ubiquitination (10) events significantly changed with AD progression in a region-specific analysis of the insoluble fraction (Table 2, adjusted p value displayed in Supp Table 2). Most modifications increased in abundance as the disease progressed, whereas a subset decreased. Several phosphorylation (pT212, pS214, pS235, pS237, pS238, pS305, pT386, pS400, pT404, pS416), acetylation (aK311, aK343, aK369), and ubiquitination (uK267, uK311, uK317) events appear to be affected early in AD, as significant changes were detected between subjects with “low” and “intermediate” AD scores. Significant regional differences in PTM abundance were observed at the “intermediate” stage (Table 2, 12 phosphorylation, 3 acetylation and 3 ubiquitination events). Among these, 6 PTMs showed progressive, stage-dependent increases, enabling clear regional stratification across AD severity (pST212, pS214, pT217, aK311, aK369 and uK343, highlighted in red in Table 2). In contrast, three events supported staging by decreasing proportionally with AD progression (pS305, pT386, and aK343, highlighted in green in Table 2). Relative abundances of all altered PTMs are illustrated in Supplementary Fig. 2.

In parallel to our primary analysis, we applied the same approach to investigate co-occurring PTMs and peptides carrying a single PTM (Supp Table 3). This approach captured, among others, canonical double-phosphorylation motifs recognized by the AT8 and PHF1 antibodies (pS202+pT205 and pS396+pS404), which are widely used in AD neuropathological assessments.

**Table 3.**
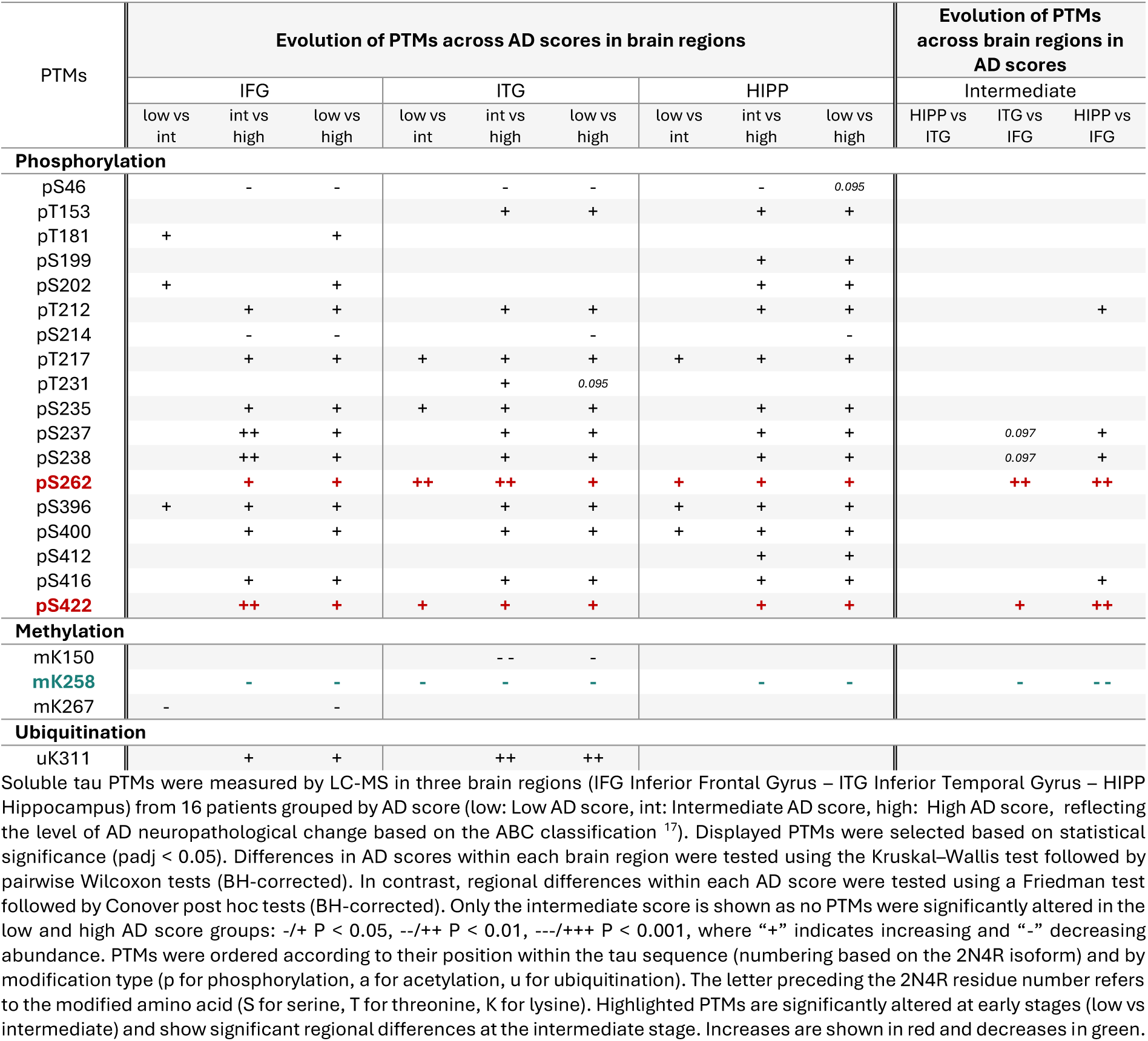
Soluble tau PTMs altered during AD progression across brain regions and between AD score.

### Soluble tau PTMs associated with tau aggregation

To better delineate the temporal biochemical sequence of tau dysregulation, we next characterized PTMs in the soluble fraction. This approach permitted the differentiation of early, potentially reversible modifications occurring on soluble tau from those associated with detergent-insoluble aggregated species that reflect established pathology.

By comparing these fractions across disease stages and brain regions, we aimed to identify PTMs that precede aggregation, thereby informing the molecular staging of tau pathology and providing potential biomarkers to track it in biological fluids. Due to the complexity of the soluble fraction, an immunoprecipitation of tau was performed before LC-MS/MS. A sequence coverage of 98% of the full-length tau protein was obtained. Across the entire cohort, we identified 42 individual tau PTMs, of which 22 were significantly associated with disease progression (Table 3 & Supp Table 4), including well-known pathology-related PTMs (e.g. pT217).

In contrast to the insoluble fraction, soluble tau displayed a more restricted pattern of PTM alterations. The phosphorylation landscape evolves gradually with disease progression, whereas ubiquitination (uK311) increases at later AD stages, consistent with the changes observed in aggregated tau. In contrast, the increase in acetylation seen in the insoluble fraction was not detected in soluble tau. Instead, we observed a progressive reduction in methylation. Canonical-phosphorylation events, including pT181, pT217 and pT231, increased with disease severity (Table 3, Fig.2 & Supp Fig.3). Notably, pS262 tracked tau pathology even more closely than the other sites previously considered. In addition, 2 methylation sites (mK258, mK267) and 8 phosphorylation sites (pT181, pS202, pT217, pS235, pS262, pS396, pS400, pS422) were significantly altered between the “low” and “intermediate” AD score in at least one brain region, highlighting their potential as early biomarkers. 6 phosphorylation and 1 ubiquitination sites showed significant regional differences (Table 3 & Supp Table 4). Notably, pS262 emerged as a robust indicator of disease progression across brain regions.

**Fig. 2.**
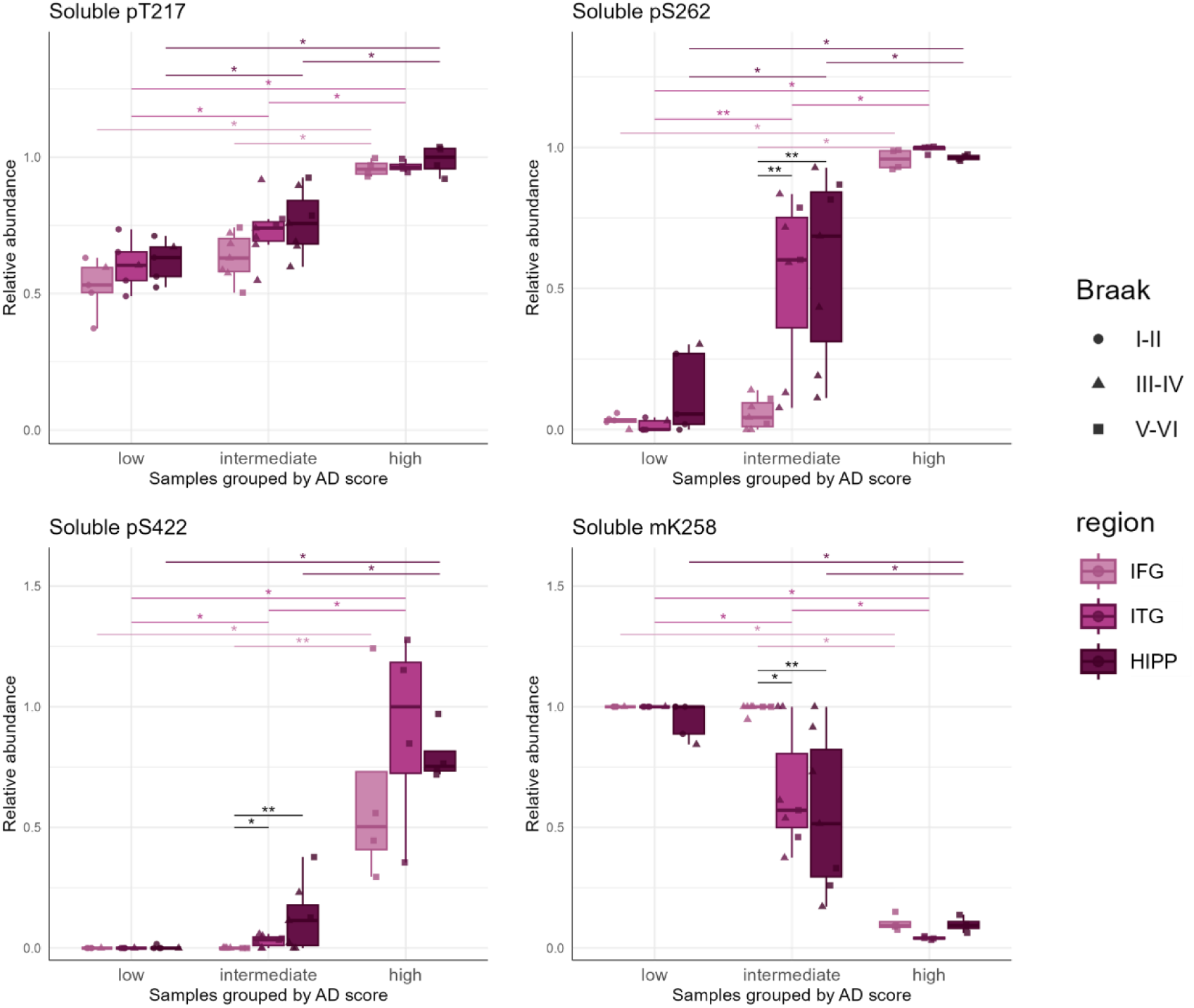
Early and progressive alterations of soluble tau PTMs across several brain regions during AD progression. Tau PTMs were measured using mass spectrometry in the soluble brain fraction of three brain regions (*IFG* Inferior Frontal Gyrus – *ITG* Inferior Temporal Gyrus – *HIPP* Hippocampus), in 16 patients grouped by AD score (low: Low AD score, int: Intermediate AD score, high: High AD score, reflecting the level of AD neuropathological change based on the ABC classification ^17^).The relative abundance corresponds to the sum of the abundances of all the peptides carrying the modification of interest, divided by the sum of the abundances of all the peptides containing this amino acid (both modified and unmodified). For visualization purposes, the median value for the highest group was normalized to 1. PTMs were selected based on the statistical analyses described in Table 3 and ordered by position within the tau sequence (numbered according to the 2N4R isoform) and by modification type (*p* for phosphorylation, *m* for methylation). The letter preceding the 2N4R residue number refers to the modified amino acid (*S* for serine, *T* for threonine, *K* for lysine). Statistical comparisons were made between AD scores for each brain region using the Wilcoxon rank-sum test after a Kruskal-Wallis test and between brain regions from patients with a given AD score using a Conover test after a Friedman test. Significant BH-adjusted p-values are indicated in black for the comparison between regions or in the color of the region for comparisons between AD scores: * > 0.05; ** > 0.01; *** > 0.001

**Fig. 3.**
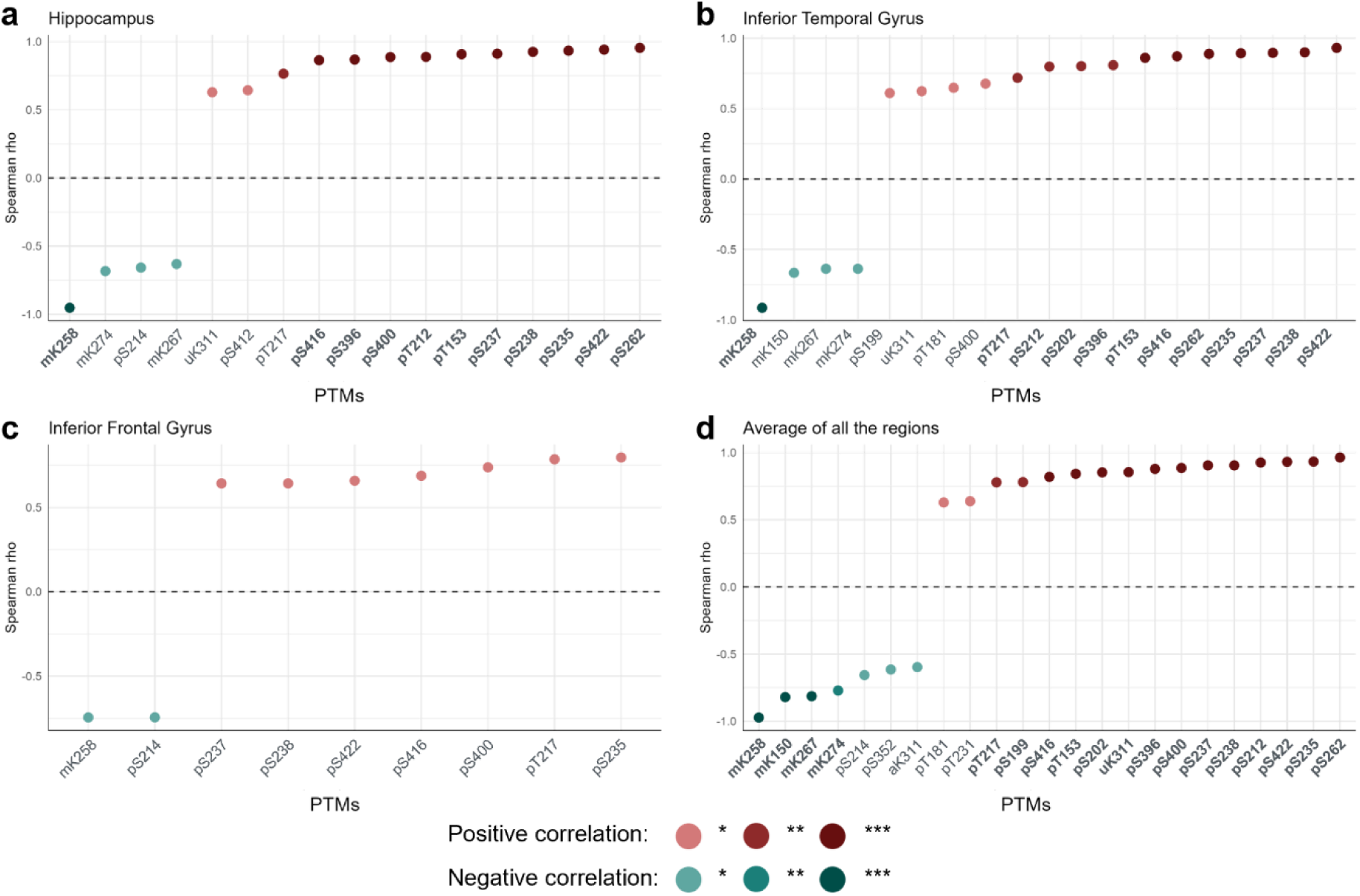
Soluble tau PTMs associated with Insoluble tau concentrations. Tau PTMs were measured using mass spectrometry in the soluble brain fractions of three brain regions (*IFG* Inferior Frontal Gyrus – *ITG* Inferior Temporal Gyrus – *HIPP* Hippocampus), in 16 subjects (spanning the spectrum of AD pathology). Insoluble tau concentration (R domains specific peptides) was measured using mass spectrometry (SureQuant method). **(a-c)** Spearman’s correlation, corrected for age, was performed for each soluble PTM in specific brain regions (p-values BH-corrected). The PTMs were aligned based on the correlation coefficient (rho). Tau PTMs numbering is based on the 2N4R isoform. The lowercase letter refers to the modification type (*p* for phosphorylation, *m* for methylation, *a* for acetylation, *u* for ubiquitination) and the uppercase letter preceding the 2N4R residue number corresponds to the modified amino acid (*S* for serine, *T* for threonine, *K* for lysine). **(d)** The same methodology was applied independently of the region, after calculating the mean abundance of each PTM for each subject. *MTBR* Microtubule binding region

As illustrated in Fig.2, distinct trajectories of PTM abundance during disease progression emerged. pT217 and pS262 exhibit a similar pattern of increase with AD progression, with a significant rise from “low” to “intermediate” and from “intermediate” to “high” in the HIPP and ITG, and elevated abundance in the IFG only at the “high” stage, consistent with the later affection of this region. Notably, pS262 enables regional staging within intermediate subjects. pS422 (and similarly pS235) displayed a lightly delayed pattern, being rarely detected in the low group and increasing thereafter. Conversely, mK258 showed a decreasing abundance with AD progression (Fig.2), similarly to other methylation sites (Supp Fig.3). Relative abundances of the other PTMs altered during AD progression are shown in Supp Fig.3.

To better define the relationship between soluble PTMs abundance and tau aggregation, we performed age-adjusted correlation analyses in each brain region. Specifically, we correlated the normalized abundance of soluble PTMs with the absolute quantification of Sarkosyl-insoluble tau (SureQuant analysis, MTBR domain) in the same region (Fig.3 a-c). In addition, we calculated the mean abundance of each PTM across the three regions to derive a global correlation with insoluble tau (Fig.3 d). Region-specific analysis identified 17, 19, and 9 soluble PTMs that were significantly correlated with insoluble tau (MTBR domain) in the HIPP, ITG, and IFG, respectively. When regional differences were not considered, 23 soluble PTMs were significantly associated with insoluble tau levels. Strikingly, methylation sites were consistently negatively correlated with tau aggregation across regions. 8 PTMs events (mK258, pT217, pS416, pS400, pS237, pS238, pS235, and pS422) correlated with tau aggregation in all three brain regions. By contrast, pS214 was significantly associated with aggregation only in HIPP and IFG. Another subset of PTMs (mK274, mK267, uK311, pS396, pT212, pT153, and pS262) was correlated with insoluble MTBR accumulation in HIPP and ITG but not in the IFG. The global analysis highlighted additional PTMs: pS352 and aK311 as negatively correlated with insoluble tau, and pT231 positively associated with tau deposition.

Taken together, these results indicate that specific soluble tau PTMs closely reflect insoluble tau accumulation and may therefore serve as robust biomarkers of aggregation. In particular, pS235, pS262 and pS422 exhibited stronger and more consistent associations with insoluble tau levels than classical fluid biomarkers such as pT181, pT217, and pT231. The systematic negative association between methylation and insoluble tau further suggests that lysine methylation may be linked to reduced aggregation propensity, raising the possibility of a modulatory or protective role in tau pathology.

In parallel, we applied the same analytical framework to examine tau PTMs detected either individually or in combination with other modifications on the same peptide (Supp Table 5 & Supp Fig.4). 26 PTMs were identified as significantly altered during AD progression in the Sarkosyl-soluble fraction (Supp Table 5). This analysis revealed that the abundance of pT231 alone decreases with AD progression, whereas its co-occurrence with nearby phosphorylation sites (such as pS235, pS237 or pS238) markedly increases. In addition, pT231 alone was strongly negatively correlated with disease severity, while pS262 and pT231+pS235 showed as strong positive association with AD progression (Supp Fig.4).

**Fig. 4.**
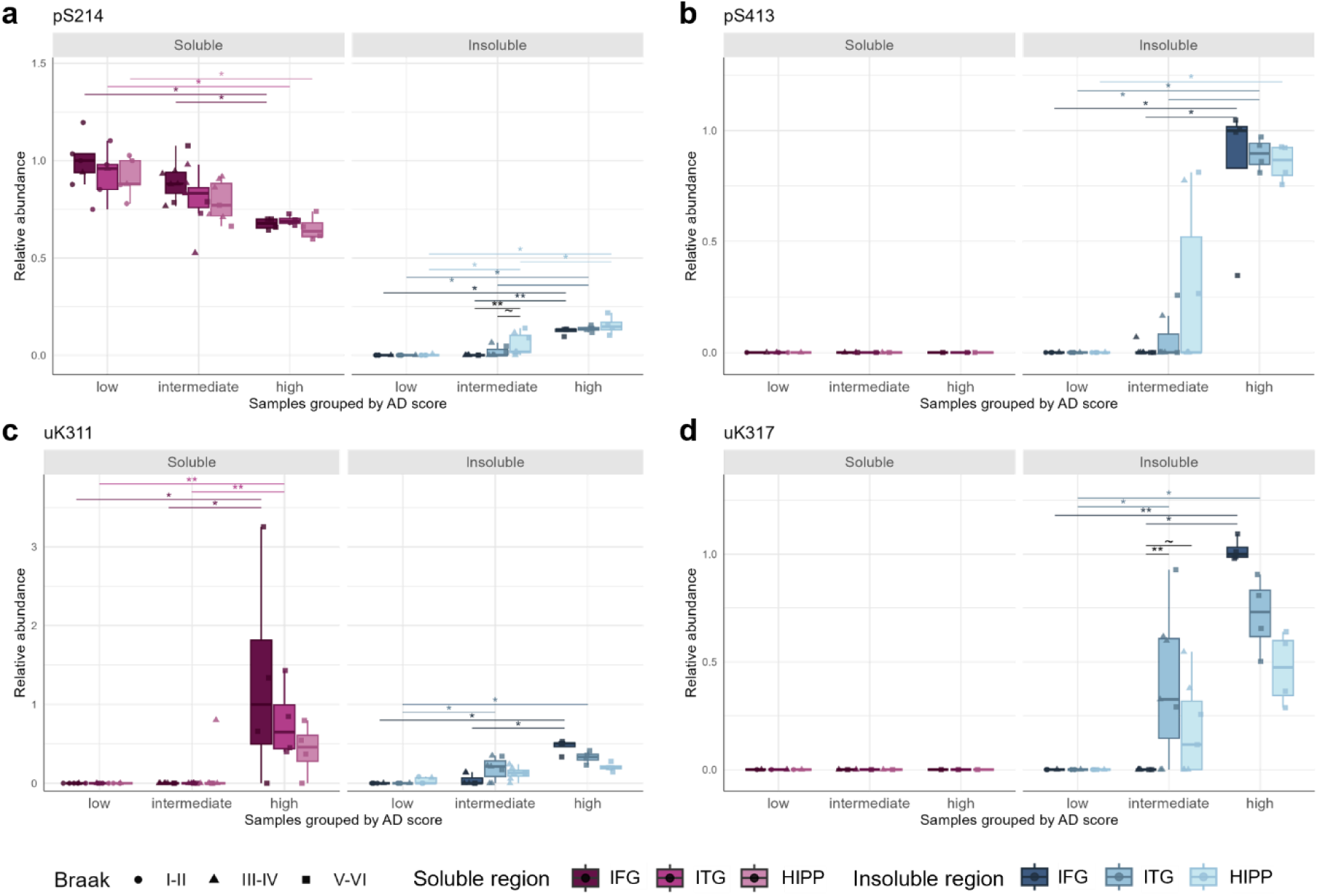
Differential tau PTMs trajectories between soluble and insoluble fractions across brain regions and AD scores. Tau PTMs were measured using mass spectrometry in the Sarkosyl-soluble and -insoluble brain fractions of three brain regions (*IFG* Inferior Frontal Gyrus – *ITG* Inferior Temporal Gyrus – *HIPP* Hippocampus), in 16 patients grouped by AD score (low: Low AD score, int: Intermediate AD score, high: High AD score, reflecting the level of AD neuropathological change based on the ABC classification ^17^).The relative abundance corresponds to the sum of the abundances of all the peptides carrying the modification of interest, divided by the sum of the abundances of all the peptides containing this amino acid (both modified and unmodified). For visualization purposes, the median value for the highest group was normalized to 1. PTMs were selected based on their differential trajectories between the two protein fractions and ordered by position within the tau sequence (numbered according to the 2N4R isoform) and by modification type (*p* for phosphorylation, *m* for methylation). The letter preceding the 2N4R residue number refers to the modified amino acid (*S* for serine, *T* for threonine, *K* for lysine). Statistical comparisons were made between AD scores for each brain region using the Wilcoxon rank-sum test after a Kruskal-Wallis test and between brain regions from patients with a given AD score using a Conover test after a Friedman test. Significant BH-adjusted p-values are indicated in black for the comparison between regions or in the color of the region for comparisons between AD scores: * > 0.05; ** > 0.01; *** > 0.001

### Differential PTM trajectories in soluble and insoluble tau during AD progression

To determine whether the identified PTMs are associated with aggregation propensity or, conversely, reflect protective mechanisms, we compared modification patterns between Sarkosyl-soluble and Sarkosyl-insoluble protein fractions across AD progressions. PTMs exhibiting the most pronounced differential trajectories between the soluble and insoluble fraction across AD scores are displayed in Fig. 4.

pS214 shows opposite trajectories in the two fractions: it increases with AD scores in the insoluble fraction but decreases in the soluble fraction. This pattern indicates a redistribution of pS214 tau toward the aggregated pool as pathology advances, supporting its association with insoluble tau accumulation. In contrast, pS413 and uK317 were identified exclusively in the insoluble fraction. Their absence from the soluble pool suggests that these modifications either occur after the onset of tau aggregation or are specifically associated with insoluble tau species, as seals of impaired processing or degradation. uK311 was detected earlier and in earlier-affected regions of the Sarkosyl-insoluble fraction than in the soluble fraction. This temporal and spatial pattern suggests that uK311 participates in early aggregation events and may mark tau tagged for degradation. Methylation events (such as mK258 and mK267) progressively decrease in abundance gradually in the soluble fraction with AD progression (Fig.2 & Supp Fig.3) and remain undetectable in the insoluble fraction. The restriction of these methylation marks to the soluble pool and their loss with disease progression support a model in which they are associated with non-aggregating tau species and may exert a protective role against aggregation.

### Overview of the dynamic changes in tau PTMs

To summarize our findings, we provide an integrated overview of the dynamic changes in tau PTMs across AD stages (Fig.5). We highlight the key PTMs that correlate with disease severity (Fig.5a). Our data indicate an early increase in phosphorylation events (pT217, pS235, pS262) accompanied by a decrease in methylation (mK258, mK267, mK274), suggesting a role for both processes in the shift towards aggregated tau species. In aggregated tau, we observed an early increase in phosphorylation, followed by a progressive increase in acetylation (aK311, aK369) and ubiquitination (uK311, uK317), suggesting a temporal sequence in which phosphorylation may accompany early aggregation. In contrast, acetylation and ubiquitination became more prominent as pathology progressed. In addition, canonical AD-related phosphorylation events were found to intensify with AD progression, reinforcing their association with pathological tau accumulation. These results are summarized in Fig. 5. In Fig. 5b and 5c, the identified significantly altered PTMs are visualized across “low”, “intermediate”, and “high” AD scores, illustrating their stage-dependent evolution in both Sarkosyl-soluble and insoluble fractions. Together, this overview links specific PTM signatures to distinct phases of tau pathology progression.

**Fig. 5.**
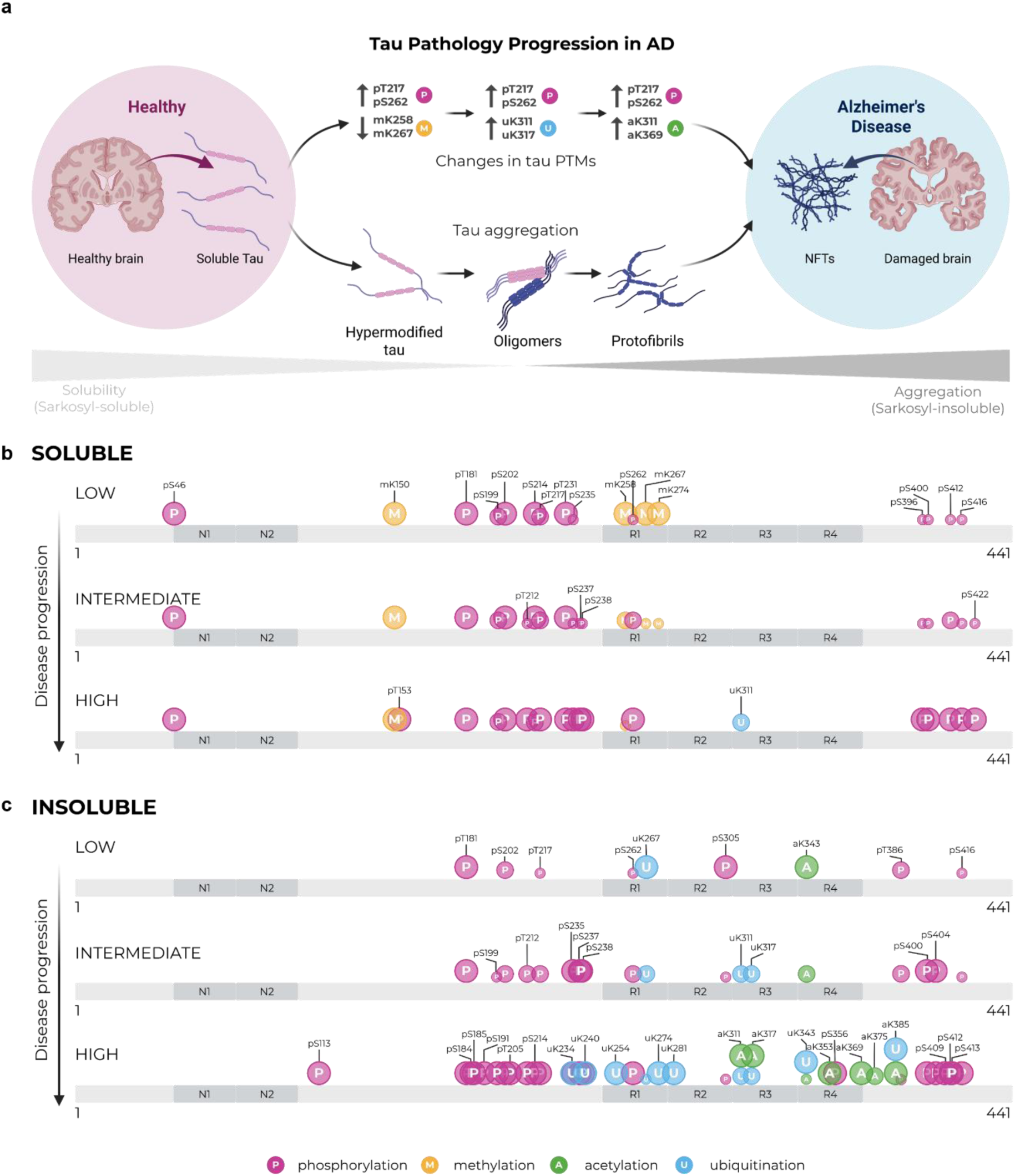
Landscape of tau PTMs during Alzheimer’s disease progression in soluble and insoluble brain fractions. **(a)** Summary of key tau PTMs during Alzheimer’s disease (AD) progression, highlighting the role of tau modifications during tau aggregation. The represented PTMs were selected based on the previous findings (Table 2&3, Supp. Table 2&4, Fig. 3). **(b-c)** Evolution of the PTMs significantly altered during AD progression, across AD score (low: Low AD score, int: Intermediate AD score, high: High AD score, reflecting the level of AD neuropathological change based on the ABC classification ^17^) in Sarkosyl-soluble and insoluble brain fractions. The represented PTMs were selected based on the statistical analysis developed in Table 2-3. All the significantly altered modifications were visualized according to their relative abundance in the Inferior Temporal Gyrus, as this region is affected at intermediate stage of AD (Braak stage III-IV). The modification type is represented using the following nomenclature: *p* for phosphorylation, *m* for methylation, *u* for ubiquitination, and *a* for acetylation. The letter preceding the 2N4R residue number refers to the modified amino acid (*S* for serine, *T* for threonine, *K* for lysine). *NFTs* neurofibrillary tangles

## DISCUSSION

Alzheimer’s disease is a multistep process in which progressive remodeling of tau PTMs accompanies and likely contributes to tau aggregation and spreading ^26,27^. Although growing attention has been devoted to tau PTMs, most biochemical studies of tau in the brain rely on post-mortem tissue from individuals at advanced stages of pathology ^25,26,33^, thereby limiting insight into early molecular events. To address this gap, we analyzed multiple brain regions from individuals with varying degrees of AD pathology, to identify diagnostic markers and molecular mechanisms underlying pathological tau deposition. In this study, we provided a regional and temporal characterization of tau PTMs during AD progression in the human brain, in both Sarkosyl-soluble and insoluble fractions. Our data reveal a staged landscape of tau modifications. Early phosphorylation events, including pT217 and pS262, emerged as promising biomarkers of disease progression, with pS262 showing the strongest association with insoluble tau accumulation. In contrast, other modifications, such as pS422 appeared to mark intermediate pathological stages, whereas uK311 was predominantly observed at advanced AD stages. Beyond their value as potential staging biomarkers, these findings suggest distinct mechanistic roles for tau modifications. Methylation appears incompatible with tau aggregation, whereas acetylation and ubiquitination are enriched in the Sarkosyl-insoluble fraction, consistent with a role of ubiquitination in tau processing and degradation that might be progressively impaired in AD conditions.

Our analysis indicates the chronology of tau PTM remodeling during AD progression. In the Sarkosyl-soluble fraction, an early hyperphosphorylation (e.g., pT217 and pS262) and hypomethylation (e.g., mK258) profile is observed. After that, PTM alterations gradually progress throughout AD, with modifications emerging at intermediate stages (e.g., pS422) and finally others at advanced stages (e.g., uK311). In the Sarkosyl-insoluble fraction, we observed progressive hyperphosphorylation of tau, along with increased acetylation (e.g., aK311 and aK369) and ubiquitination (e.g., uK311 and uK317). Of note, some modifications appear to decrease with disease progression (e.g., aK343 and uK267), suggesting dynamic remodeling even within aggregated species. Regional comparisons revealed significant differences only at the intermediate AD stage, consistent with established neuropathological staging models in which the presence of marked pathology in early-affected regions (such as the hippocampus) precedes the involvement of later-affected areas (such as the frontal lobe) ^13,17^. Importantly, our findings confirm pT217 as an early-increasing modification in AD, as previously reported and identify additional candidate markers, including pS262. This modification has been detected in the brain ^25^, both in soluble pre-tangles ^34^ and in various aggregated forms of tau ^26,33,35^. It also shows promising biomarker potential, as similar soluble pre-tangles have already been detected in the CSF ^34^. Because soluble brain tau is thought to reflect, at least partially, the composition of CSF, pS262-tau may represent a candidate fluid biomarker for early diagnosis and disease progression monitoring ^35^.

Understanding how tau PTMs regulate tau pathology is critical for developing treatments to prevent tau aggregation or improve the clearance of pathological tau. Growing evidence links AD and other neurodegenerative disorders to alterations in protein-degradation pathways, such as the proteasome and autophagy systems, highlighting the relevance of investigating PTMs involved in these processes.^36–41^ Our findings indicate a progressive increase in ubiquitination in the Sarkosyl-insoluble fraction, together with hyperphosphorylation, suggesting that ubiquitin marks insoluble tau species for degradation. In the Sarkosyl-soluble fraction, this modification appears to be delayed compared to the insoluble fraction. This observation suggests that ubiquitination marks may reflect a cellular response to pathological tau deposition, and function as signals for the clearance aggregated tau. In the context of defective protein degradation pathways (autophagy and proteasomal degradation) reported in AD^40,42,43^, ubiquitinated tau may accumulate because aggregates cannot be efficiently cleared by processes that use ubiquitination as a degradation signal. In addition, a previous study identified Otub1, a ubiquitin thioesterase, within the tau interactome. This enzyme was shown to act as a tau deubiquitinating enzyme, whose activity promotes the formation of pathological tau forms ^44^. These findings align with a broader dysregulation of ubiquitin-dependent proteostasis in AD and position ubiquitination as both a marker of advanced pathology and a readout of failed clearance mechanisms.

In contrast to phosphorylation and ubiquitination, tau methylation progressively decreased with disease progression. Although protein methylation represents a complex PTM to study in the aged brain as it increases with aging ^22^, our age-adjusted analysis identified several methylation sites in soluble tau, including mK258, that were negatively correlated with insoluble tau accumulation. Strikingly, no methylation sites were detected in the insoluble fraction, suggesting a biochemical incompatibility of lysine methylation with tau aggregation in AD. These findings are consistent with a potential protective association of tau methylation with aggregation ^45^. Mechanistically, lysine methylation may interfere with competing PTMs such as ubiquitination, thereby modulating tau solubility. This finding may also indicate the existence of a stabilizing mechanism for soluble tau, mediated by methylation perhaps and other PTMs, which is altered or lost during AD progression, thereby favoring the accumulation and aggregation of demethylated tau species. Although aggregated tau has previously been reported to be methylated ^25,46^ in a primary tauopathies ^25^ or using distinct methodological approaches ^46^, our findings indicate that, in AD, methylation is largely restricted to the soluble pool and may represent a protective biochemical state. Moreover, the decrease in methylation masking the positive charge of the lysine side-chain, together with the increase in negative charges by tau phosphorylation, may shift the protein’s isoelectric point, potentially affecting its solubility. Further mechanistic studies are required to determine if methylation directly modulates tau aggregation or merely reflects a pathological imbalance of PTMs in AD.

In the current clinical practice, preclinical AD diagnosis remains a challenge, as the clinical manifestations are often subtle or undetectable ^7,8,47^. Moreover, estimating AD progression in the brain remains challenging without costly approaches such as PET imaging, which cannot be used on a large scale. Developing fluid biomarkers that can stage AD progression is of utmost importance. This issue is particularly timely as the first disease-modifying treatments are entering the field ^48–50^. Because these therapies may work even better in preclinical stages of the disease ^9,10^, early and accurate diagnosis is critical for healthcare systems. Moreover, the ability to reliably classify patients across the AD spectrum becomes essential if therapeutic strategies are to extend beyond the earliest symptomatic stages of disease. Because biofluid tau closely resembles soluble brain tau ^29,35^, our study may provide a framework for developing fluid biomarkers that can stage AD progression. Previous studies, from our group (in a case-control model where only IFG was analyzed) and others, have investigated tau PTMs associated with AD ^25,26,28^. However, the temporal emergence of these modifications during disease progression remains poorly understood. In the present study, we reproduced most of these observations ^25,26,28^, and, more importantly, mapped the appearance and evolution of these PTMs across distinct stages of AD pathology in different brain regions. Finally, our analysis confirmed the robustness of widely studied tau biomarkers, including pT217 ^51^, while highlighting the limitations of others, such as pT231, which may rise too early in disease development and consequently shows poor correlation with aggregated tau burden in the brain.

We acknowledge that our study presents several limitations. First, the analysis was restricted to three brain regions, which may not fully reflect the spatial progression of tau PTMs across the entire brain during AD. In addition, the relatively small number of subjects per group reflects the limited availability of well-characterized post-mortem human brain tissue Nonetheless, non-parametric statistical comparisons were feasible and yielded consistent statistically significant patterns. From a technical perspective, tau immunoprecipitation had to be performed on the Sarkosyl-soluble brain fraction due to its complexity, potentially leading to an underrepresentation of certain tau species, particularly truncated forms. However, these limitations did not prevent the acquisition of high-quality mass spectrometry data spanning the entire tau sequence (98% sequence coverage), supporting the robustness of our conclusions regarding the stage-dependent remodeling of tau PTMs in AD. Finally, this study focuses on AD and does not include other tauopathies. This reflects the rarity of primary tauopathies, which limits access to well-characterized cohorts and, unlike AD, the lack of standardized staging frameworks. Nevertheless, in the context of an aging population, our findings provide valuable insights into tau pathology and its relevance to age-related neurodegeneration.

In conclusion, these findings reveal a dynamic and stage-dependent landscape of tau PTMs during AD progression: early phosphorylation changes, later ubiquitination modifications associated with tau aggregation and supporting a protein clearance defect, and decreasing tau methylation, potentially protective against tau aggregation. This work refines our understanding of the spatiotemporal evolution of tau biochemistry and provides mechanistic and translational insights into the tauopathy characteristic of AD. Future studies should assess these PTMs in biofluids to evaluate their biomarker potential, and in model systems to determine their role in tau aggregation and toxicity.

## METHODS

### Ethics

Most of the brain samples were obtained from the Netherlands Brain Bank at the Netherlands Institute for Neuroscience, Amsterdam (open access: www.brainbank.nl). All Material has been collected from donors for or from whom a written informed consent for a brain autopsy and the use of the material and clinical information for research purposes had been obtained by the NBB. The same brain samples came from the UCLouvain Brain Donation Program (UCL2020355).

Of note, one subject initially included in the “low” group (as shown in the group counts above) was excluded from our analysis because he showed marked primary age-related tauopathy (PART), which would have biased our results.

### Brain tissue processing and protein extracts preparation

Insoluble and soluble proteins were extracted from human post-mortem brain tissues. The extraction method used to separate the insoluble and soluble proteins is based on a well-established protocol ^52^ and has previously been used and validated by our laboratory ^25^. Brain pieces of 0.2g were collected by dissection while the tissue was still frozen for each subject. The dissected tissue was diced into 2 x 2 mm pieces before being homogenized in 1 mL (to reach 20 % w/v) of homogenization buffer (50 mM HEPES, pH 7, containing 250 mM sucrose and protease inhibitor cocktail (Roche #11873580001) by 20 strokes in a 2 mL Dounce homogenizer. The total homogenate (TH) volume was adjusted to 2 mL with homogenization buffer. A fraction of this TH was mixed with Sarkosyl and NaCl to final concentrations of 1% w/v and 0.2M, respectively. The Sarkosyl-homogenates (SH) were sonicated with 3 x 10 s pulses at 30% amplitude on ice using a microtip probe. 2 mL of SH were transferred to 4 mL ultracentrifuge tubes for centrifugation in a pre-cooled Beckman 50Ti fixed-angle rotor at 180,000 x g for 30 min at 4°C. The supernatants, corresponding to the Sarkosyl-soluble protein fraction (S), were recovered and collected in Lobind tubes (Eppendorf #0030108132). The pellet, corresponding to the Sarkosyl-insoluble protein fraction (P) was washed twice in cold low-saline buffer (50 mM HEPES, pH 7, 250 mM sucrose) before being solubilized by incubation in urea buffer (50 mM Tris-HCl pH 8.5, 8 M urea) for 30 min at 20°C, followed by brief sonication (1s) on ice. Protein concentration was measured using the bicinchoninic acid assay kit (Thermo Fisher Scientific #23225). Proteins were stored at -80°C.

### Immunoprecipitation and crosslink - Soluble tau enrichment

For the beads’ preparation, 37.5 µL of dynabeads (Thermo Fisher Scientific #10004D) per sample were coupled with 7.5 µg per sample of HJ8.7 antibody anti-tau antibody (D. Holtzman at Washington University School of Medicine in St Louis, Missouri, USA cat# HJ8.7, RRID:AB_2721234 [http://antibodyregistry.org/AB_2721234] ) for 2h on a rotating wheel at 4°C. Antibodies were crosslinked to the protein G using DMP in crosslink buffer (0.2M triethanolamine buffer pH 8.2, 20 mM dimethyl pimelimidate) and quenched in Tris 50 mM pH 7.5. The beads were stored in PBS containing 0.01% azide at 4°C before use to pull down tau protein from each sample. Tau proteins were enriched from the “S” fractions by immunoprecipitation (IP). For this purpose, 250 µg of the “S” fraction of each sample were incubated overnight (16h) on a rotating wheel at 4°C with the HJ8.7 crosslinked beads. The beads were washed 3 times with cold PBS to eliminate unbound proteins. The immunoprecipitated proteins were eluted in 100 µL of acid buffer (80% (v/v) acetonitrile (ACN) - 0.2% (v/v) Trifluoroacetic acid (TFA) - 0.1% (v/v) formic acid (FA)) for 20 min under strong agitation at 20°C, before being vacuum-dried. Lobind tubes (Eppendorf #0030108116) were used for all the previously described steps.

### Sample preparation for soluble tau PTMs identification (untargeted MS)

The dried post-IP eluates were resuspended in 40 µL of TEAB buffer (100 mM triethylammonium bicarbonate, pH 8.5) with 0.2 µg of sequencing-grades trypsin per sample (Promega V5111) and incubated at 37°C overnight (16h) for digestion. Digestion was stopped by adding 1 µL of 10% TFA before vacuum-drying. The peptides were resuspended in 12 µL of injection buffer (3.5% (v/v) ACN – 1.5% (v/v) TFA) before being centrifuged. 10 µL of the supernatant was transferred to MS vials and 1 µL of the mixture was injected for LC-MS/MS analyses.

### Sample preparation for insoluble tau PTMs identification (untargeted MS)

No enrichment step was performed on the insoluble fraction (P), as we previously showed that it was unnecessary to study tau PTMs in this fraction ^25^ due to its low complexity. Solubilized aggregates were diluted in 50 mM ammonium bicarbonate to reduce the urea concentration to 1M final. 4 µg of proteins were digested by incubation with 0.2 µg of sequencing grade trypsin (Promega #V5111) at 37°C for 16h. The digests were vacuum-dried and resuspended in 15 µL of injection buffer (3.5% (v/v) ACN – 1.5% (v/v) TFA) before being centrifuged. 14 µL of the supernatant was transferred to MS vials and 1.5 µL of the mixture was injected for LC-MS/MS analyses.

### Sample preparation for insoluble tau isoform absolute quantification (targeted MS)

The same methodology as for the PTMs identification in the “P” fraction was used. 4 µg of insoluble proteins were digested as described in the previous section. The peptides were resuspended in 50 mM ammonium bicarbonate, pH 9.2, for overnight agitation with 1 unit of agarose-bound alkaline phosphatase (Sigma-Aldrich #P0762) at 37°C ^25^. The samples were centrifuged at 400 x g for 4 min at 20°C. The supernatants were vacuum-dried and resuspended in 15 µL of injection buffer (3.5% (v/v) ACN – 1.5% (v/v) TFA) containing 0.5 ng/µL of AQUA-grade isotopically labelled peptides specific for each tau isoform (obtained from Synpeptide Co Ltd) before being centrifuged. 14 µL of the supernatant was transferred to MS vials, and 2 µL of the mixture was injected for LC-MS/MS analyses using SureQuant method.

### LC-MS/MS absolute quantification of tau and data analysis

MS was performed using the Orbitrap Fusion Lumos Tribrid Mass Spectrometer (Thermo Fisher Scientific). Protein digests were loaded onto a reversed-phase pre-column (Acclaim PepMap 100, Thermo Fisher Scientific, Waltham, United States) and eluted in backflush mode. Peptide separation was achieved on a reversed-phase analytical EasySpray column (Acclaim PepMap RSLC C18, 0.075 mm × 250 mm, Thermo Fisher Scientific, Waltham, United States) equilibrated in 0.1% (v/v) trifluoroacetic acid (solvent A) using a 90-min linear gradient of 0.1% (v/v) trifluoroacetic acid in 80% (v/v) acetonitrile (solvent B) at a flow rate of 300 nl/min on an Vanquish nanoHPLC system (Thermo Fisher Scientific, Waltham, United States). The Orbitrap analyser was operated in PRM mode, with a resolution of 120,000 for MS1 and 60,000 for targeted MS2 scans, an Automatic Gain Control (AGC) target of 10^6^ Ions, and maximum injection times of 50 ms in MS1 and 5×10^5^ and 10 ms in MS2. Peptides were selected for MS/MS fragmentation with a HCD collision energy of 32%, internal standard-triggered data acquisition using isolation windows of 1 Th and minimal intensities predetermined by the Thermo Fisher Scientific SureQuant method. This method enabled high-quality MS2 of target peptide ions by setting a fragmentation trigger threshold at 6 transition matches (PRM) between AQUA peptide experimental daughter ions and their theoretical counterparts. All data were analysed using Skyline softaware. Endogeneous tau quantity was calculated as on the ratio of light to heavy.

### Untargeted LC-MS/MS - PTMs identification and data analysis

Peptide samples were analyzed by nano-liquid chromatography coupled to tandem mass spectrometry (nanoLC–MS/MS). Peptides were dissolved in solvent A (0.1% TFA in 3.5% ACN) and directly loaded onto a reversed-phase pre-column (Acclaim PepMap 100, Thermo Fisher Scientific) and eluted in backflush mode. Peptide separation was performed on a C18 reversed-phase analytical column (PepMap RSLC C18, 75 μm × 250 mm, Thermo Fisher Scientific) at a flow rate of 300 nL/min using a Vanquish nanoHPLC system. Mass spectrometric analysis was performed on an Orbitrap Fusion Lumos tribrid mass spectrometer (Thermo Fisher Scientific), operated in data-dependent acquisition (DDA) mode. Full MS scans were acquired in the Orbitrap over an m/z range of 375–1800 at a resolution of 120,000, with an AGC target of 4 × 10⁵ ions and a maximum injection time of 50 ms. MS/MS spectra were acquired in the Orbitrap following HCD fragmentation at 30%. MS/MS scans were acquired with an AGC target of 1 × 10^5^ ions and a maximum injection time of 110 ms. Raw data were processed using Proteome Discoverer version 2.5 (Thermo Fisher Scientific). Protein identification was performed using the Sequest and MSAmanda search engines.

### Data management, Normalization, Statistics and Reproducibility

No statistical method was used to predetermine sample size. For all our groups and conditions, sample sizes were determined by the availability of post-mortem brain tissue in the NBB and the UCLouvain biobank. We selected the AD and cognitively normal individuals for whom hippocampus, inferior temporal gyrus, and inferior frontal gyrus were available. Moreover, 2 subjects from UCLouvain were added to the cohort as they met the same inclusion criteria. Subjects were classified into three AD scores based on the ABC score ^17^: “low” (n = 5), “intermediate” (n = 7), and “high” (n = 4). For each subject, three samples were processed, corresponding to the 3 brain regions studied. One additional “low” subject (not included in the group counts above) was excluded because he showed marked primary age-related tauopathy (PART) and was slightly older than the other individuals, which would have biased our analysis. A minimal sample size of n = 4 per group was deemed sufficient, as it allows for the detection of significant differences in tau isoforms within the aggregates. A total of n = 16 was deemed sufficient, as it allowed for the detection of correlations between PTMs and insoluble tau MTBR abundance (p < 0.05). MS data were acquired once per sample and per acquisition type. Subjects were not randomized to groups, as we are not conducting a clinical trial.

Both in targeted (tau isoforms absolute quantification) and untargeted (tau PTMs) MS data, the studied groups were compared through a multiple group comparisons Kruskal-Wallis test (p < 0.05) for the AD score comparison and through a Friedman test (p < 0.05) for the regional comparison. The pairwise group comparison procedure following the Kruskal-Wallis test was the Wilcoxon rank-sum test (p < 0.05) for the AD stage comparison. In contrast, a Conover test (p < 0.05) was performed after the Friedman test to assess the regional differences. p-values were adjusted for multiple testing using the Benjamini–Hochberg False Discovery Rate (FDR) correction, and features with an adjusted p-value (FDR) < 0.05 were considered statistically significant.

During data analysis, missing values were imputed as zero. Peptides were filtered to exclude those not consistently detected in at least one group (defined as a single brain region within a single AD score category), requiring detection in at least 25% of subjects within that group. Manual verification of peptide identifications was also performed. Despite constant amounts of total peptides across all the samples, amounts of tau peptides were variable in the insoluble fraction, as well as in the soluble fraction. Moreover, as highlighted by the absolute quantification, all tau domains are not equally represented in our samples. Therefore, we could not normalize by the total protein abundance or by a unique tau domain for all the PTMs. To ensure accurate comparisons between groups, relative abundances obtained from DDA LC-MS/MS analyses (for both soluble and insoluble protein fractions) were normalized. The relative abundance of each modification was calculated by summing the abundances of peptides carrying the modification and dividing this value by the total abundance of peptides containing the corresponding modifiable amino acid, including both modified and unmodified forms. All data processing steps were performed using Excel and RStudio. For visualization purposes, the median was set to 1 for each PTM in the figures.

## FUNDING

AV was supported by an Aspirant fellowship from the Belgian FRS-FNRS (Fonds National pour la Recherche Scientifique, 40018606/40033431). CB, MJ, and NS were funded by a Chargé de Recherche postdoctoral fellowship from the FNRS. DV was also supported by the FNRS by a CDR (research credit). This work was supported by grants from the SAO-FRA Alzheimer Research Foundation (SAO-FRA 2018/0025), UCLouvain Action de Recherche Concertée (ARC21/26-114), Fondation Louvain, Queen Elisabeth Medical Foundation (FMRE AlzHex), F.R.S.-FNRS (FNRS J.0106.22) attributed to PKC and grants from the SAO-FRA Alzheimer Research Foundation (SAO-FRA 2020/0028, SAO-FRA 2022/0028) attributed to NS. The FNRS provided grants for BH (#CCL40010417). This work was also supported by additional FNRS funding (#OL J.0099.20), for the FRFS-WELBIO under Grant n°40010035, the Queen Elizabeth Medical Foundation, the Belgian Alzheimer Research Foundation, and a Concerted Research Action (Brainbrush).

## ACKNOWLEDGEMENTS

The authors thank David Holtzman (Washington University, USA) for providing the HJ8.7 anti-tau antibody. We thank Yasmine Salman, Benoît Lengelé and Catherine Behets (UCLouvain, Belgium) for collecting and handling the UCLouvain human samples. We are grateful to Dietmar R. Thal and Sandra O. Tomé (Leuven Brain Institute, KU Leuven, Belgium) for the neuropathological assessment of these UCLouvain samples. We also thank the Netherlands Brain Bank for providing the majority of the human brain samples used in this study.

## Notes

### Competing Interest Statement

The authors have declared no competing interest.

## REFERENCES

1 Hou, Y. et al. Ageing as a risk factor for neurodegenerative disease. Nat Rev Neurol 15, 565–581 (2019). 10.1038/s41582-019-0244-7

2 Spillantini, M. G. & Goedert, M. Tau pathology and neurodegeneration. Lancet Neurol 12, 609–622 (2013). 10.1016/s1474-4422(13)70090-5

3 Gustavsson, A. et al. Global estimates on the number of persons across the Alzheimer’s disease continuum. Alzheimers Dement 19, 658–670 (2023). 10.1002/alz.12694

4 Nichols, E. et al. Estimation of the global prevalence of dementia in 2019 and forecasted prevalence in 2050: an analysis for the Global Burden of Disease Study 2019. The Lancet Public Health 7, e105–e125 (2022). 10.1016/S2468-2667(21)00249-8

5 Barthélemy, N. R. et al. Highly accurate blood test for Alzheimer’s disease is similar or superior to clinical cerebrospinal fluid tests. Nat Med 30, 1085–1095 (2024). 10.1038/s41591-024-02869-z

6 Salvadó, G. et al. Disease staging of Alzheimer’s disease using a CSF-based biomarker model. Nat Aging 4, 694–708 (2024). 10.1038/s43587-024-00599-y

7 Dubois, B., von Arnim, C. A. F., Burnie, N., Bozeat, S. & Cummings, J. Biomarkers in Alzheimer’s disease: role in early and differential diagnosis and recognition of atypical variants. Alzheimers Res Ther 15, 175 (2023). 10.1186/s13195-023-01314-6

8 Alzola, P. et al. Neuropsychological Assessment for Early Detection and Diagnosis of Dementia: Current Knowledge and New Insights. J Clin Med 13 (2024). 10.3390/jcm13123442

9 Zhang, Y., Chen, H., Li, R., Sterling, K. & Song, W. Amyloid β-based therapy for Alzheimer’s disease: challenges, successes and future. Signal Transduct Target Ther 8, 248 (2023). 10.1038/s41392-023-01484-7

10 Jucker, M. & Walker, L. C. Alzheimer’s disease: From immunotherapy to immunoprevention. Cell 186, 4260–4270 (2023). 10.1016/j.cell.2023.08.021

11 Lebouvier, T., Pasquier, F. & Buée, L. Update on tauopathies. Curr Opin Neurol 30, 589–598 (2017). 10.1097/wco.0000000000000502

12 Arendt, T., Stieler, J. T. & Holzer, M. Tau and tauopathies. Brain Res Bull 126, 238–292 (2016). 10.1016/j.brainresbull.2016.08.018

13 Braak, H. & Braak, E. Neuropathological stageing of Alzheimer-related changes. Acta Neuropathol 82, 239–259 (1991). 10.1007/bf00308809

14 Hardy, J. A. & Higgins, G. A. Alzheimer’s disease: the amyloid cascade hypothesis. Science 256, 184–185 (1992). 10.1126/science.1566067

15 Frisoni, G. B. et al. The probabilistic model of Alzheimer disease: the amyloid hypothesis revised. Nat Rev Neurosci 23, 53–66 (2022). 10.1038/s41583-021-00533-w

16 Kepp, K. P., Robakis, N. K., Høilund-Carlsen, P. F., Sensi, S. L. & Vissel, B. The amyloid cascade hypothesis: an updated critical review. Brain 146, 3969–3990 (2023). 10.1093/brain/awad159

17 Hyman, B. T. et al. National Institute on Aging-Alzheimer’s Association guidelines for the neuropathologic assessment of Alzheimer’s disease. Alzheimers Dement 8, 1–13 (2012). 10.1016/j.jalz.2011.10.007

18 DeTure, M. A. & Dickson, D. W. The neuropathological diagnosis of Alzheimer’s disease. Mol Neurodegener 14, 32 (2019). 10.1186/s13024-019-0333-5

19 Thal, D. R., Rüb, U., Orantes, M. & Braak, H. Phases of A beta-deposition in the human brain and its relevance for the development of AD. Neurology 58, 1791–1800 (2002). 10.1212/wnl.58.12.1791

20 Mirra, S. S. et al. The Consortium to Establish a Registry for Alzheimer’s Disease (CERAD). Part II. Standardization of the neuropathologic assessment of Alzheimer’s disease. Neurology 41, 479–486 (1991). 10.1212/wnl.41.4.479

21 Braak, H., Alafuzoff, I., Arzberger, T., Kretzschmar, H. & Del Tredici, K. Staging of Alzheimer disease-associated neurofibrillary pathology using paraffin sections and immunocytochemistry. Acta Neuropathol 112, 389–404 (2006). 10.1007/s00401-006-0127-z

22 Alquezar, C., Arya, S. & Kao, A. W. Tau Post-translational Modifications: Dynamic Transformers of Tau Function, Degradation, and Aggregation. Front Neurol 11, 595532 (2020). 10.3389/fneur.2020.595532

23 Guillozet-Bongaarts, A. L. et al. Phosphorylation and cleavage of tau in non-AD tauopathies. Acta Neuropathol 113, 513–520 (2007). 10.1007/s00401-007-0209-6

24 Iqbal, K. et al. Tau pathology in Alzheimer disease and other tauopathies. Biochim Biophys Acta 1739, 198–210 (2005). 10.1016/j.bbadis.2004.09.008

25 Kyalu Ngoie Zola, N., et al. Specific post-translational modifications of soluble tau protein distinguishes Alzheimer’s disease and primary tauopathies. Nat Commun 14, 3706 (2023). 10.1038/s41467-023-39328-1

26 Wesseling, H. et al. Tau PTM Profiles Identify Patient Heterogeneity and Stages of Alzheimer’s Disease. Cell 183, 1699–1713.e1613 (2020). 10.1016/j.cell.2020.10.029

27 Arakhamia, T. et al. Posttranslational Modifications Mediate the Structural Diversity of Tauopathy Strains. Cell 180, 633–644.e612 (2020). 10.1016/j.cell.2020.01.027

28 Kametani, F. et al. Comparison of Common and Disease-Specific Post-translational Modifications of Pathological Tau Associated With a Wide Range of Tauopathies. Front Neurosci 14, 581936 (2020). 10.3389/fnins.2020.581936

29 Han, P. et al. A Quantitative Analysis of Brain Soluble Tau and the Tau Secretion Factor. J Neuropathol Exp Neurol 76, 44–51 (2017). 10.1093/jnen/nlw105

30 Horie, K., Barthélemy, N. R., Sato, C. & Bateman, R. J. CSF tau microtubule binding region identifies tau tangle and clinical stages of Alzheimer’s disease. Brain 144, 515–527 (2021). 10.1093/brain/awaa373

31 Li, L. et al. Alzheimer’s disease brain contains tau fractions with differential prion-like activities. Acta Neuropathol Commun 9, 28 (2021). 10.1186/s40478-021-01127-4

32 Aragão Gomes, L., et al. Maturation of neuronal AD-tau pathology involves site-specific phosphorylation of cytoplasmic and synaptic tau preceding conformational change and fibril formation. Acta Neuropathol 141, 173–192 (2021). 10.1007/s00401-020-02251-6

33 Kumar, M. et al. Alzheimer proteopathic tau seeds are biochemically a forme fruste of mature paired helical filaments. Brain 147, 637–648 (2024). 10.1093/brain/awad378

34 Islam, T. et al. Phospho-tau serine-262 and serine-356 as biomarkers of pre-tangle soluble tau assemblies in Alzheimer’s disease. Nat Med 31, 574–588 (2025). 10.1038/s41591-024-03400-0

35 Horie, K. et al. Regional correlation of biochemical measures of amyloid and tau phosphorylation in the brain. Acta Neuropathol Commun 8, 149 (2020). 10.1186/s40478-020-01019-z

36 Hara, T. et al. Suppression of basal autophagy in neural cells causes neurodegenerative disease in mice. Nature 441, 885–889 (2006). 10.1038/nature04724

37 Komatsu, M. et al. Loss of autophagy in the central nervous system causes neurodegeneration in mice. Nature 441, 880–884 (2006). 10.1038/nature04723

38 Puangmalai, N. et al. Lysine 63-linked ubiquitination of tau oligomers contributes to the pathogenesis of Alzheimer’s disease. J Biol Chem 298, 101766 (2022). 10.1016/j.jbc.2022.101766

39 Schmidt, M. F., Gan, Z. Y., Komander, D. & Dewson, G. Ubiquitin signalling in neurodegeneration: mechanisms and therapeutic opportunities. Cell Death Differ 28, 570–590 (2021). 10.1038/s41418-020-00706-7

40 Zhang, W., Xu, C., Sun, J., Shen, H. M., Wang, J. & Yang, C. Impairment of the autophagy-lysosomal pathway in Alzheimer’s diseases: Pathogenic mechanisms and therapeutic potential. Acta Pharm Sin B 12, 1019–1040 (2022). 10.1016/j.apsb.2022.01.008

41 Wang, L., Sooram, B., Kumar, R., Schedin-Weiss, S., Tjernberg, L. O. & Winblad, B. Tau degradation in Alzheimer’s disease: Mechanisms and therapeutic opportunities. Alzheimers Dement 21, e70048 (2025). 10.1002/alz.70048

42 Jiang, S. et al. Early proteasome downregulation and dysfunction drive proteostasis failure in Alzheimer’s disease. Brain 148, 4372–4388 (2025). 10.1093/brain/awaf222

43 Piras, A., Collin, L., Grüninger, F., Graff, C. & Rönnbäck, A. Autophagic and lysosomal defects in human tauopathies: analysis of post-mortem brain from patients with familial Alzheimer disease, corticobasal degeneration and progressive supranuclear palsy. Acta Neuropathol Commun 4, 22 (2016). 10.1186/s40478-016-0292-9

44 Wang, P. et al. Tau interactome mapping based identification of Otub1 as Tau deubiquitinase involved in accumulation of pathological Tau forms in vitro and in vivo. Acta Neuropathol 133, 731–749 (2017). 10.1007/s00401-016-1663-9

45 Balmik, A. A. & Chinnathambi, S. Methylation as a key regulator of Tau aggregation and neuronal health in Alzheimer’s disease. Cell Commun Signal 19, 51 (2021). 10.1186/s12964-021-00732-z

46 Thomas, S. N. et al. Dual modification of Alzheimer’s disease PHF-tau protein by lysine methylation and ubiquitylation: a mass spectrometry approach. Acta Neuropathol 123, 105–117 (2012). 10.1007/s00401-011-0893-0

47 Mankhong, S. et al. Development of Alzheimer’s Disease Biomarkers: From CSF- to Blood-Based Biomarkers. Biomedicines 10 (2022). 10.3390/biomedicines10040850

48 Sims, J. R. et al. Donanemab in Early Symptomatic Alzheimer Disease: The TRAILBLAZER-ALZ 2 Randomized Clinical Trial. Jama 330, 512–527 (2023). 10.1001/jama.2023.13239

49 van Dyck, C. H. et al. Lecanemab in Early Alzheimer’s Disease. N Engl J Med 388, 9–21 (2023). 10.1056/NEJMoa2212948

50 Rafii, M. S. & Aisen, P. S. Detection and treatment of Alzheimer’s disease in its preclinical stage. Nat Aging 3, 520–531 (2023). 10.1038/s43587-023-00410-4

51 Warmenhoven, N. et al. A comprehensive head-to-head comparison of key plasma phosphorylated tau 217 biomarker tests. Brain 148, 416–431 (2025). 10.1093/brain/awae346

52 Diner, I., Nguyen, T. & Seyfried, N. T. Enrichment of Detergent-insoluble Protein Aggregates from Human Postmortem Brain. JoVE, e55835 (2017). doi:10.3791/55835

