## Supplementary Information for "Stage-dependent tau post-translational modifications map the spatiotemporal progression of Alzheimer’s disease"

### Supplementary Table1 | Subjects and sample characteristics

| Subject | Origin | Age at death | Sex | ApoE | Brain Weight | PMD | Diagnosis at death | Thal | Braak | ABC code | AD score |
|---|---|---|---|---|---|---|---|---|---|---|---|
| *A* | UCL | 90 | M | | | | NC | 0 | I | A0B1C0 | low |
| *B* | NBB | 84 | M | | 1150 | 320 | NC | 1 | I | A1B1C0 | low |
| *C* | NBB | 89 | M | | 1200 | 575 | NC | 1 | II | A1B1C0 | low |
| *D* | NBB | 99 | F | 32 | 1100 | 245 | NC | 1 | II | A1B1C0 | low |
| *E* | NBB | 87 | M | 33 | 1265 | 385 | NC | 1 | III | A1B2C0 | low |
| **F* | NBB | 102 | F | 33 | 1093 | 310 | NC | 1 | IV | A1B2C0 | low |
| *G* | NBB | 92 | M | 43 | 1360 | 445 | NC | 3 | III | A2B2C1 | intermediate |
| *H* | NBB | 96 | F | | 1020 | 265 | NC | 3 | III | A2B2C1 | intermediate |
| *I* | NBB | 92 | M | | 1395 | 490 | NC | 3 | III | A2B2C2 | intermediate |
| *J* | NBB | 92 | F | 33 | 1090 | 350 | NC | 4 | IV | A3B2C1 | intermediate |
| *K* | NBB | 85 | M | | 1375 | 490 | AD | 3 | IV | A2B2C2 | intermediate |
| *L* | NBB | 94 | F | | 1013 | 350 | AD | 4 | V | A3B2C2 | intermediate |
| *M* | NBB | 98 | F | 43 | 1090 | 365 | AD | 4 | V | A3B3C2 | intermediate |
| *N* | NBB | 64 | M | | 1450 | 298 | AD | 4 | V | A3B3C3 | high |
| *O* | NBB | 70 | F | | 885 | 375 | AD | 5 | VI | A3B3C3 | high |
| *P* | NBB | 74 | M | | 925 | 325 | AD | 5 | VI | A3B3C3 | high |
| *Q* | UCL | 79 | M | | | 1440 | AD | 5 | VI | A3B3C3 | high |

Characteristics of 17 subjects studied: origin (*NBB* Netherlands Brain Bank or *UCL* Université Catholique de Louvain Belgium), age at death (years), sex (*M* for male or *F* for female), ApoE status, brain weight (g), PMD *post mortem* delay (minutes between death and autopsy), diagnosis at death (*NC* non-demented control or *AD* Alzheimer's Disease), Thal Phases (from 0 to 5), Braak Stages (from I to VI), ABC score (from A0B0C0 to A3B3C3) and AD score (staging used in this study, adapted from on the neuropathological ABC score[1]). The *F subject was excluded from our analysis due to a marked primary age-related tauopathy.

**Supplementary Figure 1 | Quantification of insoluble tau concentration during Alzheimer's Disease progression across several brain regions**

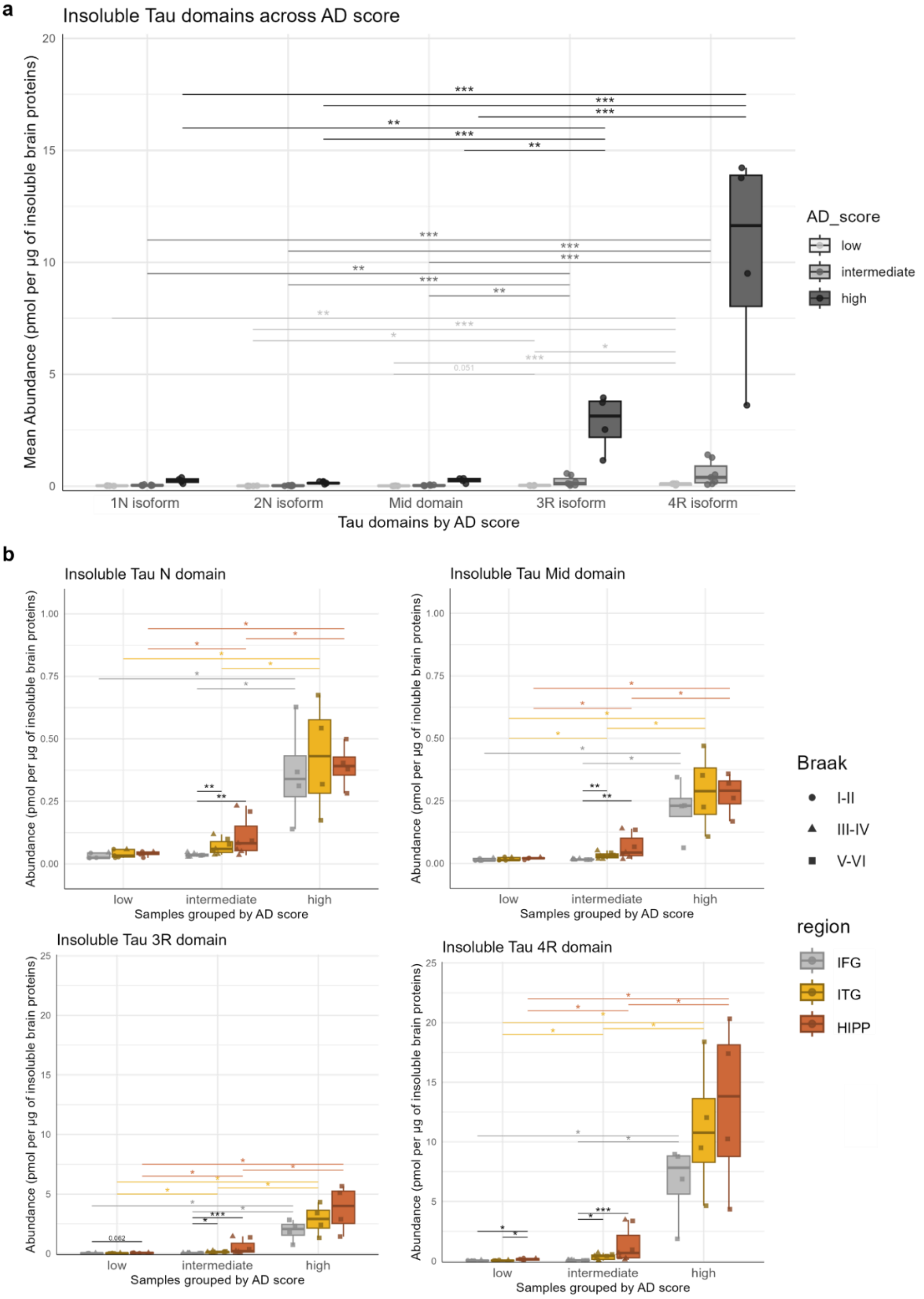


**Supp. Fig1 | Quantification of insoluble tau concentration during Alzheimer's Disease progression across several brain regions**. Tau domains and isoforms concentration from insoluble protein fraction was measured using mass spectrometry (SureQuant method) in three brain regions (IFG Inferior Frontal Gyrus – ITG Inferior Temporal Gyrus – HIPP Hippocampus) from 16 patients grouped by AD score (low: Low AD score, int: Intermediate AD score, high: High AD score, based on the neuropathological ABC score[1]). **(a)** Mean of abundance of tau domains/isoforms was calculated for each subject. Statistical comparisons were made to assess the abundance of each domain/isoform within specific AD scores using a Conover test. Significant BH-adjusted p-values are indicated in the color of the AD scores: * > 0.05; ** > 0.01; *** > 0.001 **(b)** Each tau domains/isoforms concentration were studied during Alzheimer's Disease (AD) progression. Statistical comparisons were made between AD scores for each brain region using the Wilcoxon rank-sum test after a Kruskal-Wallis test and between brain regions from patients with a given AD score using a Conover test after a Friedman test. Significant BH-adjusted p-values are indicated in black for the comparison between regions or in the color of the region for comparisons between AD scores: * > 0.05; ** > 0.01; *** > 0.001

### Supplementary Table 2 | Insoluble tau PTMs altered during AD progression across brain regions and between AD score

| PTMs | Evolution of PTMs across AD scores in brain regions | | | | | | | | | | | | Evolution of PTMs across brain regions in AD scores | | | | | | Change with AD |
|---|---|---|---|---|---|---|---|---|---|---|---|---|---|---|---|---|---|---|---|
| | **Kruskal-Wallis test** | | | IFG | | | ITG | | | HIPP | | | **Friedman test** | | | Intermediate | | | |
| | **IFG** | **ITG** | **HIPP** | low vs int | int vs high | low vs high | low vs int | int vs high | low vs high | low vs int | int vs high | low vs high | **Low** | **Inter.** | **High** | Hipp vs ITG | ITG vs IFG | Hipp vs IFG | |
| **Phosphorylation** | | | | | | | | | | | | | | | | | | | |
| pS113 | 0.010 | 0.014 | 0.031 | | 0.009 | | | | 0.032 | | | 0.032 | | | | | | | + |
| pT181 | 0.033 | | | | 0.018 | | | | | | | | | 0.021 | | | | 0.031 | + |
| pS184 | 0.010 | 0.004 | 0.045 | | 0.032 | 0.042 | | 0.006 | 0.011 | | | 0.032 | | | | | | | + |
| pS185 | 0.010 | 0.004 | 0.045 | | 0.032 | 0.042 | | 0.006 | 0.011 | | | 0.032 | | | | | | | + |
| pS191 | 0.003 | 0.015 | | | 0.006 | 0.011 | | | 0.032 | | | | | | | | | | + |
| pS199 | 0.003 | 0.010 | 0.031 | | 0.006 | 0.011 | | 0.023 | 0.023 | | | 0.045 | | 0.074 | | | 0.038 | 0.032 | + |
| pS202 | 0.019 | 0.017 | | | 0.024 | 0.024 | | 0.024 | 0.024 | | | | | 0.024 | | | | | + |
| pT205 | 0.010 | 0.004 | 0.019 | | 0.032 | 0.042 | | 0.006 | 0.011 | | 0.016 | 0.016 | | | | | | | + |
| pT212 | 0.010 | 0.007 | 0.028 | | 0.016 | 0.016 | 0.012 | 0.012 | 0.012 | | 0.024 | 0.024 | | 0.018 | | 0.026 | 0.008 | 0.000 | + |
| pS214 | 0.003 | 0.009 | 0.019 | | 0.006 | 0.011 | | 0.016 | 0.016 | 0.024 | 0.024 | 0.024 | | 0.024 | | 0.058 | | 0.002 | + |
| pT217 | 0.015 | 0.011 | 0.028 | | 0.018 | 0.024 | | 0.018 | 0.024 | | | 0.048 | | 0.018 | | 0.070 | 0.024 | 0.001 | + |
| pS235 | 0.023 | 0.011 | | 0.038 | | 0.038 | 0.016 | | 0.027 | | | | | | | | | | + |
| pS237 | 0.011 | 0.035 | | | 0.046 | 0.032 | 0.045 | | 0.032 | | | | | | | | | | + |
| pS238 | 0.007 | 0.014 | | | 0.016 | 0.016 | 0.045 | | 0.032 | | | | | 0.024 | | | 0.036 | 0.003 | + |
| pS262 | 0.015 | 0.011 | 0.031 | | 0.018 | 0.024 | | 0.018 | 0.024 | | | 0.048 | | 0.074 | | | | 0.063 | + |
| pS305 | 0.015 | 0.014 | 0.036 | | 0.018 | 0.024 | | 0.018 | 0.024 | | | 0.048 | | 0.028 | | 0.059 | 0.008 | 0.000 | - |
| pS356 | | 0.004 | 0.027 | | | | | 0.006 | 0.011 | | | 0.032 | | | | | | | + |
| pT386 | 0.010 | 0.014 | 0.025 | | 0.018 | 0.024 | | 0.018 | 0.024 | 0.024 | | 0.024 | | 0.021 | | 0.009 | 0.009 | 0.000 | - |
| pS400 | 0.010 | 0.012 | | 0.029 | 0.018 | 0.023 | 0.018 | | 0.018 | | | | | 0.055 | | | 0.006 | 0.003 | + |
| pT404 | 0.031 | 0.027 | | | | 0.032 | 0.045 | | 0.032 | | | | | | | | | | + |
| pS409 | 0.003 | 0.009 | 0.019 | | 0.006 | 0.011 | | 0.016 | 0.016 | | 0.023 | 0.023 | | 0.032 | | | | 0.017 | + |
| pS412 | 0.003 | 0.009 | 0.019 | | 0.006 | 0.011 | | 0.016 | 0.016 | | 0.023 | 0.023 | | 0.032 | | | | 0.017 | + |
| pS413 | 0.007 | 0.009 | 0.027 | | 0.016 | 0.016 | | 0.016 | 0.016 | | | 0.032 | | | | | | | + |
| pS416 | 0.011 | 0.013 | 0.019 | | 0.027 | 0.027 | | 0.024 | 0.024 | 0.009 | 0.009 | 0.016 | | | | | | | + |
| **Acetylation** | | | | | | | | | | | | | | | | | | | |
| aK311 | 0.003 | 0.007 | 0.019 | | 0.006 | 0.011 | | 0.016 | 0.016 | 0.030 | 0.030 | 0.030 | | 0.032 | | 0.004 | | 0.002 | + |
| aK317 | 0.010 | 0.004 | | | 0.032 | 0.042 | | 0.006 | 0.011 | | | | | | | | | | + |
| aK343 | 0.012 | 0.007 | | | 0.018 | 0.024 | 0.009 | 0.009 | 0.016 | | | | | 0.018 | | 0.003 | 0.014 | 0.000 | - |
| aK353 | 0.003 | 0.004 | | | 0.006 | 0.011 | | 0.006 | 0.011 | | | | | | | | | | + |
| aK369 | 0.003 | 0.009 | 0.019 | | 0.006 | 0.011 | | 0.016 | 0.016 | 0.030 | 0.027 | 0.027 | | 0.084 | | 0.041 | | 0.007 | + |
| aK375 | 0.011 | 0.004 | 0.031 | | 0.046 | 0.032 | | 0.006 | 0.011 | | | 0.032 | | | | | | | + |
| aK385 | 0.003 | 0.007 | 0.022 | | 0.006 | 0.011 | | 0.016 | 0.016 | | 0.034 | 0.032 | | | | | | | + |
| **Ubiquitination** | | | | | | | | | | | | | | | | | | | |
| uK234 | 0.010 | 0.004 | 0.022 | | 0.032 | 0.042 | | 0.006 | 0.011 | | 0.034 | 0.032 | | | | | | | + |
| uK240 | 0.047 | 0.011 | 0.019 | | | | | 0.032 | 0.042 | | 0.027 | 0.027 | | | | | | | + |
| uK254 | 0.010 | 0.039 | 0.031 | | 0.032 | 0.042 | | | 0.032 | | | 0.032 | | | | | | | + |
| uK267 | 0.010 | 0.010 | 0.019 | | 0.018 | 0.024 | 0.048 | 0.018 | 0.024 | 0.018 | 0.018 | 0.018 | | 0.038 | | | 0.017 | 0.014 | - |
| uK274 | 0.010 | 0.011 | 0.031 | | 0.032 | 0.042 | | 0.022 | 0.023 | | | 0.032 | | | | | | | + |
| uK281 | 0.010 | 0.010 | 0.019 | | 0.032 | 0.042 | | 0.027 | 0.027 | | 0.016 | 0.016 | | | | | | | + |
| uK311 | 0.010 | 0.013 | 0.028 | | 0.016 | 0.016 | 0.045 | | 0.032 | | | | | | | | | | + |
| uK317 | 0.003 | 0.016 | 0.028 | | 0.006 | 0.011 | 0.045 | | 0.032 | | | 0.032 | | 0.061 | | | 0.009 | 0.075 | + |
| uK343 | 0.010 | 0.004 | 0.045 | | 0.032 | 0.042 | | 0.006 | 0.011 | | 0.045 | 0.032 | | 0.029 | | 0.000 | | 0.000 | + |
| uK385 | 0.047 | 0.011 | 0.019 | | | | | 0.032 | 0.042 | | 0.027 | 0.027 | | | | | | | + |

Insoluble tau PTMs were measured by mass spectrometry in three brain regions (IFG Inferior Frontal Gyrus – ITG Inferior Temporal Gyrus – HIPP Hippocampus) from 16 patients grouped by AD score (low: Low AD score, int: Intermediate AD score, high: High AD score based on the neuropathological ABC score [1]). Displayed PTMs were selected based on statistical significance (padj < 0.05). Differences in AD scores within each brain region were tested using the Kruskal–Wallis test followed by pairwise Wilcoxon tests (BH-corrected). In contrast, regional differences within each AD score were tested using a Friedman test followed by Conover post hoc tests (BH-corrected). Only the intermediate score is shown as no PTMs were significantly altered in the low and high AD score groups. The “Change with AD” column reflects the direction of variation in modification abundance across AD progression. “+” indicates increasing abundance and “-” decreasing abundance. PTMs were ordered according to their position within the tau sequence (numbering based on the 2N4R isoform) and by modification type (*p* for phosphorylation, *a* for acetylation, *u* for ubiquitination). The letter preceding the 2N4R residue number refers to the modified amino acid (*S* for serine, *T* for threonine, *K* for lysine).

**Supplementary Figure 2 | Insoluble tau PTMs altered during AD progression across brain regions and between AD score**

**a Phosphorylation**

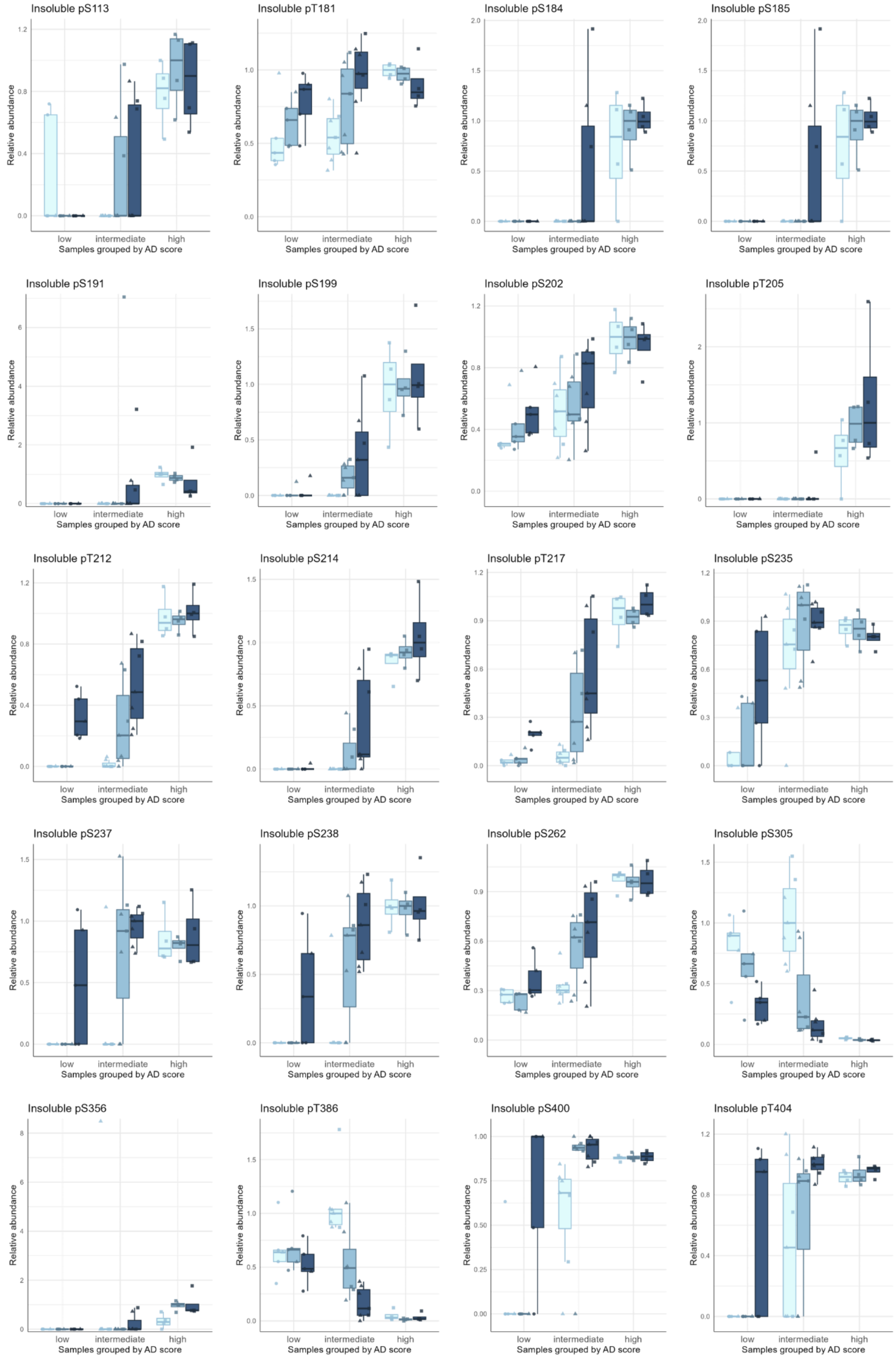

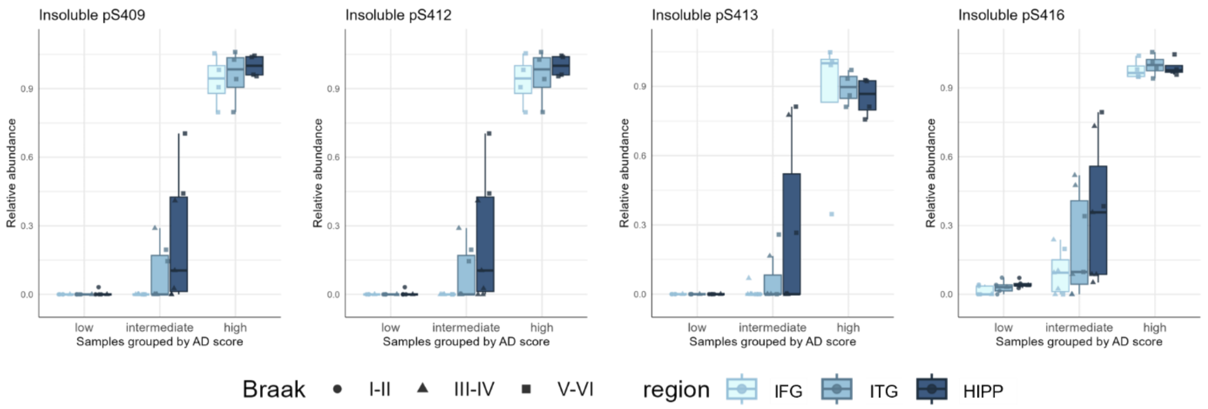


**b Ubiquitination**

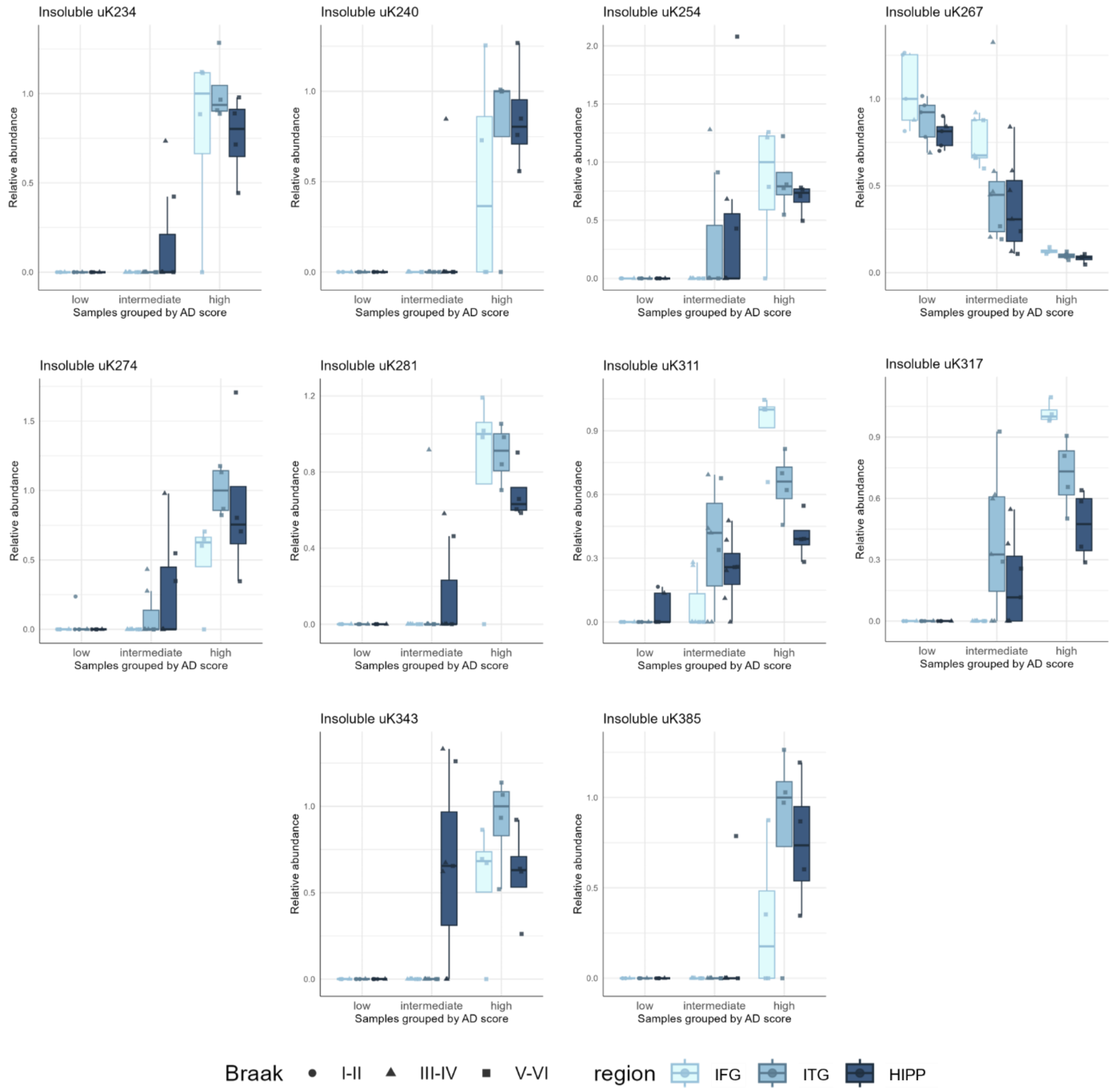

**c Acetylation**

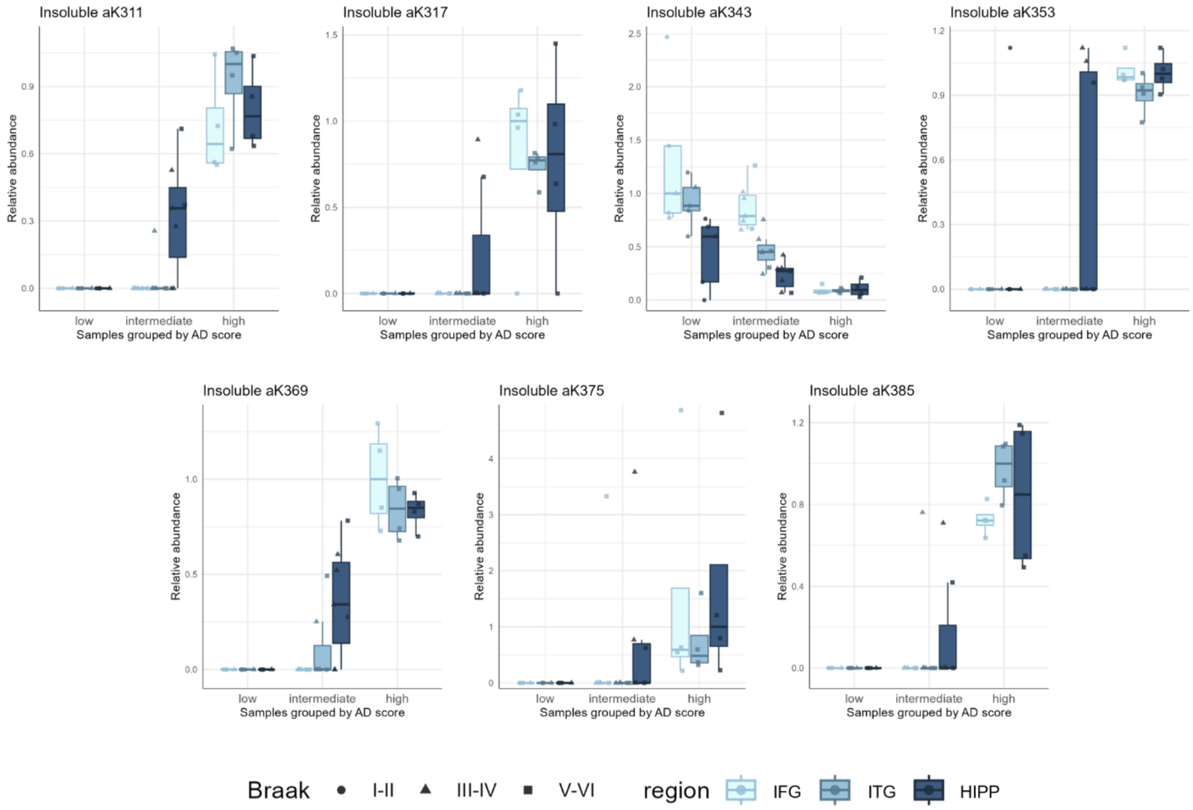


**Supp. Fig. 2 | Insoluble tau PTMs altered during AD progression across brain regions and between AD score regions.** Tau PTMs were measured using mass spectrometry in the insoluble brain fraction of three brain regions (*IFG* Inferior Frontal Gyrus – *ITG* Inferior Temporal Gyrus – *HIPP* Hippocampus), in 16 patients grouped by AD score (*low*: Low AD score, *int*: Intermediate AD score, *high*: High AD score, based on the neuropathological ABC score [1]). The relative abundance corresponds to the sum of the abundances of all the peptides carrying the modification of interest, divided by the sum of the abundances of all the peptides containing this amino acid (both modified and unmodified). For visualization purposes, the highest median value among all groups was normalized to 1. All the significantly altered PTMs were displayed (statistical analysis from Supp. Table 2) and ordered according to their position within the tau sequence (numbering based on the 2N4R isoform) and by modification type (p for phosphorylation, m for methylation). The letter preceding the 2N4R residue number refers to the modified amino acid (*S* for serine, *T* for threonine, *K* for lysine). Statistical comparisons were made between AD scores for each brain region using the Wilcoxon rank-sum test after a Kruskal-Wallis test, and between brain regions from patients with a given AD score using the Conover test after a Friedman test. Significant BH-adjusted p-values are indicated in black for the comparison between regions or in the color of the region for comparisons between AD scores: * > 0.05; ** > 0.01; *** > 0.001

### Supplementary Table 3 | Insoluble tau PTMs combinations altered during AD progression across brain regions and between AD score

| PTMs | Evolution of PTMs across AD scores in brain regions | | | | | | | | | | | | Evolution of PTMs across AD scores in brain regions | | | | | | Change with AD |
|---|---|---|---|---|---|---|---|---|---|---|---|---|---|---|---|---|---|---|---|
| | Kruskal-Wallis test | | | IFG | | | ITG | | | HIPP | | | Friedman test | | | Intermediate | | | |
| | IFG | ITG | HIPP | low vs int | int vs high | low vs high | low vs int | int vs high | low vs high | low vs int | int vs high | low vs high | low | inter | high | hipp vs ITG | ITG vs IFG | Hipp vs IFG | |
| pS113 | | 0.012 | 0.037 | | | | | 0.034 | 0.032 | | | 0.032 | | | | | | | + |
| pT181 | | | | | | | | | | | | | | 0.034 | | | | 0.042 | + |
| pT181+pS184+pS185 | 0.010 | 0.003 | 0.042 | | 0.032 | 0.042 | | 0.006 | 0.011 | | | 0.032 | | | | | | | + |
| pT181+pS191 | 0.010 | 0.003 | | | 0.032 | 0.042 | | 0.006 | 0.011 | | | | | | | | | | + |
| pS191 | 0.004 | 0.017 | | | 0.006 | 0.011 | | | 0.032 | | | | | | | | | | + |
| pS199+pS202+pT205 | 0.010 | 0.003 | 0.020 | | 0.032 | 0.042 | | 0.006 | 0.011 | | 0.016 | 0.016 | | | | | | | + |
| pT212+pS214+pT217 | 0.004 | 0.010 | 0.020 | | 0.006 | 0.011 | | 0.016 | 0.016 | 0.024 | 0.024 | 0.024 | | 0.034 | | 0.058 | 0.080 | 0.002 | + |
| pT217 | 0.016 | 0.012 | 0.030 | | 0.018 | 0.024 | | 0.018 | 0.024 | 0.027 | | 0.027 | | 0.025 | | | 0.065 | 0.005 | + |
| pT231 | 0.015 | | 0.037 | | 0.018 | 0.024 | | | | 0.045 | | 0.045 | | 0.066 | | | 0.041 | 0.041 | - |
| uK234+pT231+pS235 | 0.010 | 0.003 | 0.023 | | 0.032 | 0.042 | | 0.006 | 0.011 | | 0.034 | 0.032 | | | | | | | + |
| pT231+pS235 | 0.029 | | | | | 0.045 | | | | | | | | | | | | | + |
| pT231+pS235+pS238 | 0.004 | | | | 0.006 | 0.011 | | | | | | | | | | | | | + |
| uK254 | 0.010 | 0.046 | 0.034 | | 0.032 | 0.042 | | | 0.032 | | | 0.032 | | | | | | | + |
| pS262 | 0.015 | 0.012 | 0.042 | | 0.018 | 0.024 | | 0.018 | 0.024 | | | 0.048 | | 0.088 | | | | 0.063 | + |
| uK267+pS262 | 0.010 | | 0.042 | | 0.032 | 0.042 | | | | | | 0.032 | | | | | | | + |
| uK267 | 0.010 | 0.011 | 0.020 | | 0.018 | 0.024 | 0.048 | 0.018 | 0.024 | 0.018 | 0.018 | 0.018 | | 0.047 | | | 0.017 | 0.014 | - |
| uK274 | 0.010 | 0.012 | 0.034 | | 0.032 | 0.042 | | 0.022 | 0.023 | | | 0.032 | | | | | | | + |
| uK281 | 0.010 | 0.011 | 0.020 | | 0.032 | 0.042 | | 0.027 | 0.027 | | 0.016 | 0.016 | | | | | | | + |
| pS305 | 0.016 | 0.015 | 0.037 | | 0.018 | 0.024 | | 0.018 | 0.024 | | | 0.048 | | 0.034 | | 0.080 | 0.007 | 0.000 | - |
| aK311 | 0.004 | 0.006 | 0.020 | | 0.006 | 0.011 | | 0.016 | 0.016 | 0.030 | 0.030 | 0.030 | | 0.040 | | | 0.004 | 0.002 | + |
| uK311 | 0.010 | 0.014 | 0.030 | | 0.016 | 0.016 | 0.045 | | 0.032 | | | | | | | | | | + |
| uK317 | 0.004 | 0.018 | 0.030 | | 0.006 | 0.011 | 0.045 | | 0.032 | | | 0.032 | | 0.072 | | | 0.009 | 0.075 | + |
| aK317 | 0.010 | 0.003 | | | 0.032 | 0.042 | | 0.006 | 0.011 | | | | | | | | | | + |
| pS324 | | | 0.020 | | | | | | | 0.013 | | 0.016 | | | | | | | - |
| aK343 | 0.013 | 0.006 | | | 0.018 | 0.024 | 0.009 | 0.009 | 0.016 | | | | | 0.025 | | 0.003 | 0.001 | 0.000 | - |
| uK343 | 0.010 | 0.003 | 0.043 | | 0.032 | 0.042 | | 0.006 | 0.011 | 0.045 | | 0.032 | | 0.035 | | 0.000 | | 0.000 | + |
| aK353 | 0.004 | 0.003 | | | 0.006 | 0.011 | | 0.006 | 0.011 | | | | | | | | | | + |
| pS356 | | 0.003 | 0.029 | | | | | 0.006 | 0.011 | | | 0.032 | | | | | | | + |
| aK375 | 0.012 | 0.003 | 0.034 | | 0.046 | 0.032 | | 0.006 | 0.011 | | | 0.032 | | | | | | | + |
| aK385 | 0.004 | 0.006 | 0.023 | | 0.006 | 0.011 | | 0.016 | 0.016 | | 0.034 | 0.032 | | | | | | | + |
| uK385 | 0.049 | 0.012 | 0.020 | | | | | 0.032 | 0.042 | | 0.027 | 0.027 | | | | | | | + |
| pT386 | 0.010 | 0.015 | 0.027 | | 0.018 | 0.024 | | 0.018 | 0.024 | 0.024 | | 0.024 | | 0.025 | | 0.009 | 0.009 | 0.000 | - |
| pS396 | 0.015 | 0.020 | 0.048 | | 0.018 | 0.024 | | 0.024 | 0.024 | | | 0.045 | | | | | | | + |
| pS396+pS400+pS404 | 0.010 | 0.009 | 0.020 | | 0.016 | 0.016 | 0.030 | 0.016 | 0.016 | 0.017 | 0.018 | 0.019 | | 0.025 | | | 0.034 | 0.003 | + |
| pS396+pS400 | 0.010 | 0.012 | 0.031 | | 0.016 | 0.016 | 0.045 | 0.047 | 0.032 | | | | | 0.089 | | | 0.066 | 0.045 | + |
| uK395+pS396+pS400+pS404 | 0.010 | 0.003 | 0.020 | | 0.032 | 0.042 | | 0.006 | 0.011 | | 0.016 | 0.016 | | | | | | | + |
| pS409+pS412+pT416 | 0.004 | 0.010 | 0.020 | | 0.006 | 0.011 | | 0.016 | 0.016 | | 0.023 | 0.023 | | 0.040 | | | | 0.017 | + |
| pT416 | 0.049 | | 0.030 | | | | | | | 0.024 | | 0.024 | | | | | | | + |

Insoluble tau PTMs (detected alone or combine with others) were measured by mass spectrometry in three brain regions (*IFG* Inferior Frontal Gyrus – *ITG* Inferior Temporal Gyrus – *HIPP* Hippocampus) from 16 patients grouped by AD score (*low*: Low AD score, *int*: Intermediate AD score, *high*: High AD score, based on the neuropathological ABC score [1]). Displayed PTMs were selected based on statistical significance (padj < 0.05). Differences in AD scores within each brain region were tested using the Kruskal–Wallis test followed by pairwise Wilcoxon tests (BH-corrected). In contrast, regional differences within each AD score were tested using a Friedman test followed by Conover post hoc tests (BH-corrected). Only the intermediate score is shown as no PTMs were significantly altered in the low and high AD score groups. The "Change with AD" column reflects the direction of variation in modification abundance across AD progression. "+" indicates increasing abundance and "-" decreasing abundance. PTMs were ordered according to their position within the tau sequence (numbering based on the 2N4R isoform) and by modification type (*p* for phosphorylation, *a* for acetylation, *u* for ubiquitination). The letter preceding the 2N4R residue number refers to the modified amino acid (*S* for serine, *T* for threonine, *K* for lysine).

### Supplementary Table 4 | Soluble tau PTMs altered during AD progression across brain regions and between AD score

| PTMs | Evolution of PTMs across AD scores in brain regions | | | | | | | | | | | | Evolution of PTMs across brain regions in AD scores | | | | | | Change with AD |
|---|---|---|---|---|---|---|---|---|---|---|---|---|---|---|---|---|---|---|---|
| | Kruskal-Wallis test | | | IFG | | | ITG | | | HIPP | | | Friedman test | | | Intermediate | | | |
| | IFG | ITG | HIPP | low vs int | int vs high | low vs high | low vs int | int vs high | low vs high | low vs int | int vs high | low vs high | Low | Inter. | High | Hipp vs ITG | ITG vs IFG | Hipp vs IFG | |
| **Phosphorylation** | | | | | | | | | | | | | | | | | | | |
| pS46 | 0.032 | 0.033 | 0.048 | | 0.018 | 0.048 | | 0.018 | 0.024 | | 0.018 | | | | | | | | - |
| pT153 | | 0.014 | 0.022 | | | | | 0.016 | 0.016 | | 0.016 | 0.016 | | | | | | | + |
| pT181 | 0.032 | | | 0.024 | | 0.024 | | | | | | | | | | | | | + |
| pS199 | | | 0.036 | | | | | | | | 0.018 | 0.024 | | | | | | | + |
| pS202 | 0.030 | 0.047 | 0.036 | 0.027 | | 0.027 | | | | | 0.029 | 0.029 | | | | | | | + |
| pT212 | 0.015 | 0.019 | 0.036 | | 0.016 | 0.016 | | 0.023 | 0.023 | | 0.044 | 0.044 | | 0.043 | | | | 0.034 | + |
| pS214 | 0.030 | 0.047 | 0.036 | | 0.018 | 0.024 | | | 0.048 | | | 0.048 | | | | | | | - |
| pT217 | 0.030 | 0.016 | 0.022 | | 0.018 | 0.024 | 0.048 | 0.018 | 0.024 | 0.048 | 0.024 | 0.024 | | 0.089 | | | | | + |
| pT231 | | 0.047 | | | | | | 0.018 | | | | | | | | | | | + |
| pS235 | 0.031 | 0.014 | 0.022 | | 0.018 | 0.024 | 0.015 | 0.015 | 0.016 | | 0.018 | 0.024 | | 0.043 | | | | | + |
| pS237 | 0.009 | 0.014 | 0.022 | | 0.006 | 0.011 | | 0.016 | 0.016 | | 0.016 | 0.016 | | 0.075 | | | 0.097 | 0.039 | + |
| pS238 | 0.009 | 0.014 | 0.022 | | 0.006 | 0.011 | | 0.016 | 0.016 | | 0.016 | 0.016 | | 0.075 | | | 0.097 | 0.039 | + |
| pS262 | 0.031 | 0.014 | 0.022 | | 0.024 | 0.024 | 0.009 | 0.009 | 0.018 | 0.030 | 0.018 | 0.024 | | 0.039 | | | 0.003 | 0.001 | + |
| pS396 | 0.030 | 0.016 | 0.022 | 0.048 | 0.018 | 0.024 | | 0.018 | 0.024 | 0.048 | 0.018 | 0.024 | | | | | | | + |
| pS400 | 0.030 | 0.027 | 0.022 | | 0.018 | 0.024 | | 0.018 | 0.024 | 0.030 | 0.018 | 0.024 | | | | | | | + |
| pS412 | | | 0.036 | | | | | | | | 0.024 | 0.024 | | | | | | | + |
| pS416 | 0.030 | 0.016 | 0.022 | | 0.018 | 0.011 | | 0.018 | 0.024 | | 0.018 | 0.024 | | 0.039 | | | | 0.042 | + |
| pS422 | 0.009 | 0.014 | 0.022 | | 0.006 | | 0.030 | 0.016 | 0.016 | | 0.023 | 0.023 | | 0.043 | | | 0.029 | 0.007 | + |
| **Methylation** | | | | | | | | | | | | | | | | | | | |
| mK150 | | 0.014 | 0.042 | | | | | 0.006 | 0.011 | | | | | | | | | | - |
| mK258 | 0.015 | 0.014 | 0.022 | | 0.016 | 0.016 | 0.030 | 0.016 | 0.016 | | 0.018 | 0.027 | | 0.043 | | | 0.023 | 0.006 | - |
| mK267 | 0.031 | 0.047 | | 0.031 | | 0.031 | | | | | | | | | | | | | - |
| mK274 | 0.041 | 0.047 | | | | | | | | | | | | | | | | | - |
| **Ubiquitination** | | | | | | | | | | | | | | | | | | | |
| uK311 | 0.030 | 0.014 | | | 0.032 | 0.042 | | 0.006 | 0.011 | | | | | | | | | | + |

Soluble tau PTMs were measured by mass spectrometry in three brain regions (IFG Inferior Frontal Gyrus – ITG Inferior Temporal Gyrus – HIPP Hippocampus) from 16 patients grouped by AD score (low: Low AD score, int: Intermediate AD score, high: High AD score, based on the neuropathological ABC score [1]). Displayed PTMs were selected based on statistical significance (padj < 0.05). Differences in AD scores within each brain region were tested using the Kruskal–Wallis test followed by pairwise Wilcoxon tests (BH-corrected). In contrast, regional differences within each AD score were tested using a Friedman test followed by Conover post hoc tests (BH-corrected). Only the intermediate score is shown as no PTMs were significantly altered in the low and high AD score groups. The "Change with AD" column reflects the direction of variation in modification abundance across AD progression. "+" indicates increasing abundance and "-" decreasing abundance. PTMs were ordered according to their position within the tau sequence (numbering based on the 2N4R isoform) and by modification type (*p* for phosphorylation, *a* for acetylation, *u* for ubiquitination). The letter preceding the 2N4R residue number refers to the modified amino acid (*S* for serine, *T* for threonine, *K* for lysine).

**Supplementary Figure 3 | Soluble tau PTMs altered during AD progression across brain regions and between AD scores**

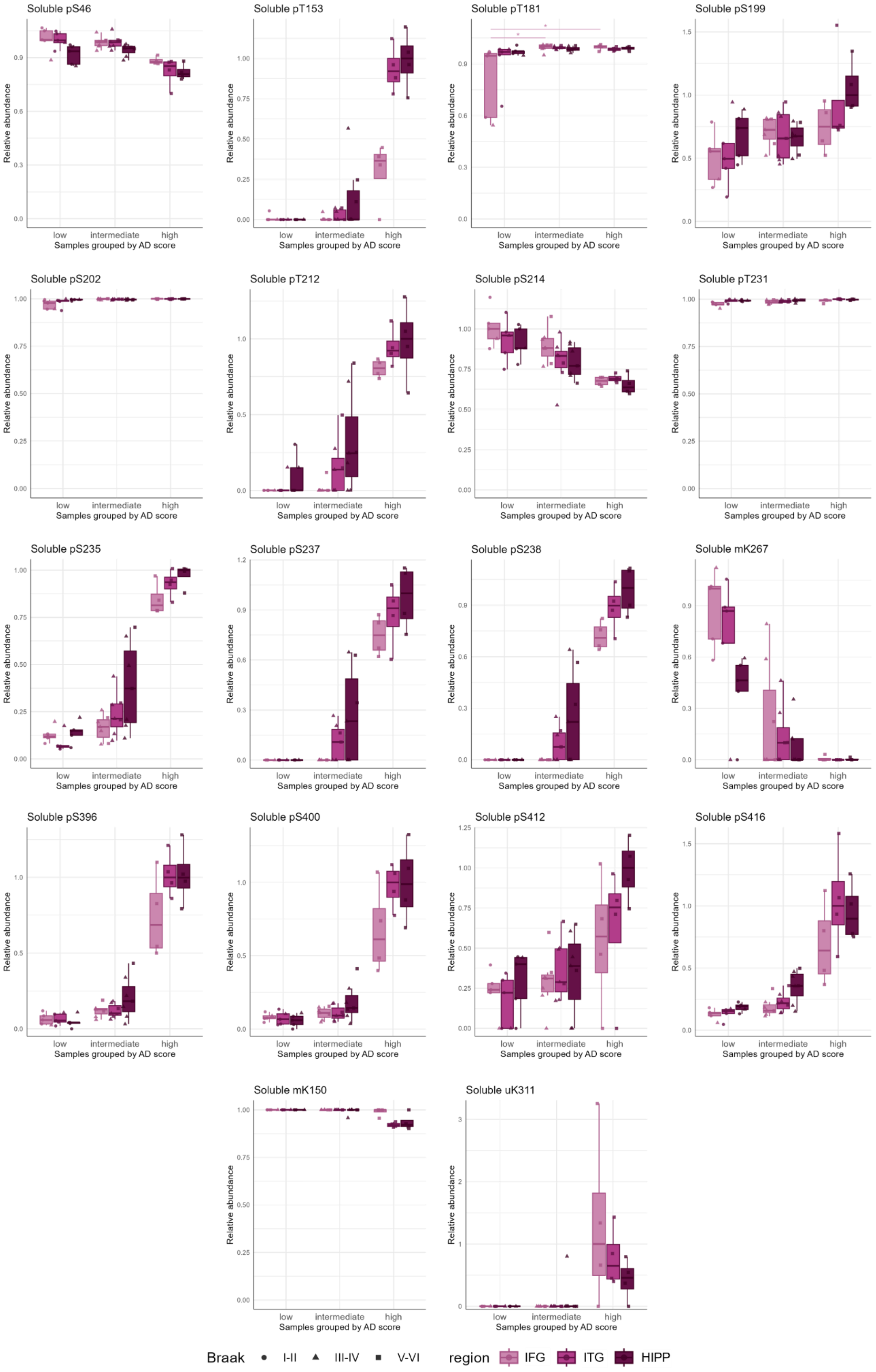

**Supp. Fig. 3 | Soluble tau PTMs altered during AD progression across brain regions and between AD scores.** Tau PTMs were measured using mass spectrometry in the soluble brain fraction of three brain regions (*IFG* Inferior Frontal Gyrus – *ITG* Inferior Temporal Gyrus – *HIPP* Hippocampus), in 16 patients grouped by AD score (*low*: Low AD score, *int*: Intermediate AD score, *high*: High AD score, based on the neuropathological ABC score [1]). The relative abundance corresponds to the sum of the abundances of all the peptides carrying the modification of interest, divided by the sum of the abundances of all the peptides containing this amino acid (both modified and unmodified). For visualization purposes, the highest median value among all groups was normalized to 1. All the significantly altered PTMs were displayed (statistical analysis from Supp. Table 4) and ordered according to their position within the tau sequence (numbering based on the 2N4R isoform) and by modification type (p for phosphorylation, m for methylation). The letter preceding the 2N4R residue number refers to the modified amino acid (*S* for serine, *T* for threonine, *K* for lysine). Statistical comparisons were made between AD scores for each brain region using the Wilcoxon rank-sum test after a Kruskal-Wallis test, and between brain regions from patients with a given AD score using the Conover test after a Friedman test. Significant BH-adjusted p-values are indicated in black for the comparison between regions or in the color of the region for comparisons between AD scores: * > 0.05; ** > 0.01; *** > 0.001. The 4 others significantly altered PTMs in the soluble fraction are displayed in Fig. 2.

### Supplementary Table 5 | Soluble tau PTMs combinations altered during AD progression across brain regions and between AD score

| PTMs | Evolution of PTMs across AD scores in brain regions | | | | | | | | | | | | Evolution of PTMs across AD scores in brain regions | | | | | | Change with AD |
|---|---|---|---|---|---|---|---|---|---|---|---|---|---|---|---|---|---|---|---|
| | **Kruskal-Wallis test** | | | IFG | | | ITG | | | HIPP | | | **Friedman test** | | | Intermediate | | | |
| | **IFG** | **ITG** | **HIPP** | low vs int | int vs high | low vs high | low vs int | int vs high | low vs high | low vs int | int vs high | low vs high | **low** | **inter** | **high** | hipp vs ITG | ITG vs IFG | Hipp vs IFG | |
| pT39+pT50 | 0.032 | 0.012 | 0.031 | | 0.018 | 0.048 | 0.015 | 0.015 | | | 0.036 | | | 0.080 | | 0.006 | | 0.061 | - |
| aK44 | | 0.030 | | | | | | 0.044 | 0.044 | | | | | | | | | | + |
| pS46 | 0.032 | 0.012 | 0.048 | | 0.018 | 0.048 | 0.048 | 0.018 | 0.024 | | 0.036 | 0.048 | | | | | | | - |
| mK150 | | 0.012 | 0.036 | | | | | 0.006 | 0.011 | | | | | | | | | | - |
| pT153 | | 0.012 | 0.017 | | | | | 0.016 | 0.016 | | 0.016 | 0.016 | | | | | | | + |
| pT175+pT181 | | | 0.036 | | | | | | | | 0.024 | 0.024 | | | | | | | + |
| pT181 | 0.042 | | | 0.030 | | 0.048 | | | | | | | | | | | | | + |
| pS199+pS202 | | | 0.031 | | | | | | | | 0.018 | 0.024 | | | | | | | + |
| pT217 | 0.032 | | 0.036 | | 0.018 | 0.024 | | | | | | 0.048 | | 0.080 | | | | | + |
| pT212+pS214+pT217 | 0.018 | 0.015 | 0.031 | | 0.016 | 0.016 | | 0.023 | 0.023 | | 0.044 | 0.044 | | 0.048 | | | | 0.034 | + |
| pS214 | 0.029 | 0.012 | 0.017 | | 0.018 | 0.024 | 0.048 | 0.018 | 0.024 | 0.048 | 0.024 | 0.024 | | 0.080 | | | | | - |
| pS214+pT217 | 0.029 | 0.017 | 0.017 | | 0.023 | 0.023 | | 0.027 | 0.027 | | 0.027 | 0.027 | | | | | | | + |
| pT231 | 0.032 | 0.012 | 0.017 | | 0.018 | 0.024 | 0.015 | 0.015 | 0.016 | | 0.018 | 0.024 | | 0.045 | | | | | - |
| pT231+pS235 | 0.029 | 0.012 | 0.017 | | 0.018 | 0.024 | 0.018 | 0.018 | 0.018 | | 0.018 | 0.024 | | 0.036 | | | | 0.064 | + |
| pT231+pS235+pS238 | 0.009 | 0.012 | 0.017 | | 0.006 | 0.011 | | 0.016 | 0.016 | | 0.016 | 0.016 | | 0.080 | | | | 0.039 | + |
| pT231+pS237+pS238 | 0.009 | 0.012 | 0.017 | | 0.006 | 0.011 | | 0.016 | 0.016 | | 0.016 | 0.016 | | 0.080 | | | 0.097 | 0.039 | + |
| pS262 | 0.032 | 0.012 | 0.017 | | 0.024 | 0.024 | 0.009 | 0.009 | 0.018 | 0.030 | 0.018 | 0.024 | | 0.038 | | | 0.003 | 0.001 | + |
| mK258 | 0.029 | 0.022 | 0.017 | | 0.022 | | | 0.029 | 0.029 | 0.050 | 0.018 | 0.029 | | | | | | | - |
| uK311 | 0.029 | 0.012 | | | 0.032 | 0.042 | | 0.006 | 0.011 | | | | | | | | | | + |
| pS356 | | 0.012 | | | | | | 0.016 | 0.016 | | | | | | | | | | + |
| pS396+pS400 | 0.029 | 0.012 | 0.017 | 0.048 | 0.018 | 0.024 | | 0.018 | 0.024 | 0.018 | 0.018 | 0.018 | | | | | | | + |
| pS396+pS400+pS404 | 0.029 | 0.012 | 0.017 | | 0.032 | 0.042 | | 0.016 | 0.023 | | 0.016 | 0.016 | | | | | | | + |
| pS396+pS404 | 0.029 | 0.012 | 0.017 | | 0.018 | 0.024 | 0.048 | 0.018 | 0.024 | 0.048 | 0.018 | 0.024 | | | | | | | + |
| pS412 | | | 0.031 | | | | | | | | 0.024 | 0.024 | | | | | | | + |
| pS416+pS422 | 0.009 | 0.012 | 0.017 | | 0.006 | 0.011 | 0.030 | 0.016 | 0.016 | | 0.023 | 0.023 | | 0.045 | | | 0.029 | 0.007 | + |
| pS416 | 0.032 | 0.012 | 0.017 | | 0.036 | 0.036 | | 0.018 | 0.024 | | 0.018 | 0.024 | | 0.060 | | | | 0.046 | + |

Soluble tau PTMs (detected alone or combine with others) were measured by mass spectrometry in three brain regions (*IFG* Inferior Frontal Gyrus – *ITG* Inferior Temporal Gyrus – *HIPP* Hippocampus) from 16 patients grouped by AD score (*low*: Low AD score, *int*: Intermediate AD score, *high*: High AD score, based on the neuropathological ABC score [1]). Displayed PTMs were selected based on statistical significance (padj < 0.05). Differences in AD scores within each brain region were tested using the Kruskal–Wallis test followed by pairwise Wilcoxon tests (BH-corrected). In contrast, regional differences within each AD score were tested using a Friedman test followed by Conover post hoc tests (BH-corrected). Only the intermediate score is shown as no PTMs were significantly altered in the low and high AD score groups. The “Change with AD” column reflects the direction of variation in modification abundance over the course of AD progression. “+” indicates increasing abundance and “-” decreasing abundance. PTMs were ordered according to their position within the tau sequence (numbering based on the 2N4R isoform) and by modification type (*p* for phosphorylation, *a* for acetylation, *u* for ubiquitination). The letter preceding the 2N4R residue number refers to the modified amino acid (*S* for serine, *T* for threonine, *K* for lysine).

**Supplementary Figure 4 | Soluble tau PTMs combinations associated with Insoluble tau concentrations.**

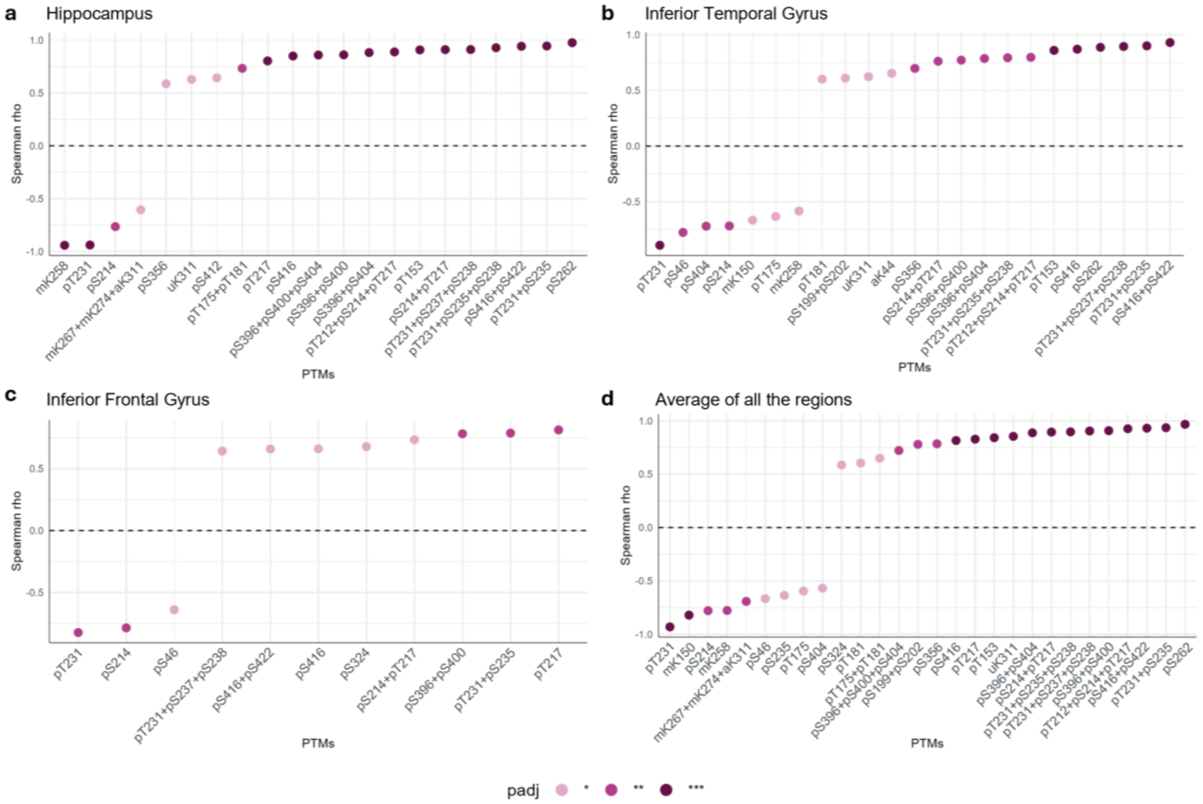


**Supp. Fig. 4 | Soluble tau PTMs combinations associated with Insoluble tau concentrations.** Tau PTMs (detected alone on their peptide or detected combined with others) were measured using mass spectrometry in the soluble brain fractions of three brain regions (*IFG* Inferior Frontal Gyrus – *ITG* Inferior Temporal Gyrus – *HIPP* Hippocampus), in 16 subjects (spanning the spectrum of AD pathology). Insoluble tau concentration (R domains specific peptides) was measured using mass spectrometry (SureQuant method). **(a-c)** Spearman's correlation, corrected for age, was performed for each soluble PTM (alone or combined on their peptides) in specific brain regions (BH-corrected p-values). The PTMs were aligned based on the correlation coefficient (rho). Tau PTMs numbering is based on the 2N4R isoform. The lowercase letter refers to the modification type (*p* for phosphorylation, *m* for methylation, *a* for acetylation, *u* for ubiquitination) and the uppercase letter preceding the 2N4R residue number corresponds to the modified amino acid (*S* for serine, *T* for threonine, *K* for lysine). **(d)** The same methodology was applied independently of the region, after calculating the mean abundance of each PTM (alone or combined on their peptides) per subject.
*MTBR* Microtubule binding region

1 Hyman, B. T. *et al.* National Institute on Aging-Alzheimer's Association guidelines for the neuropathologic assessment of Alzheimer's disease. *Alzheimers Dement* **8**, 1-13 (2012). https://doi.org:10.1016/j.jalz.2011.10.007
